# PRC1 drives Polycomb-mediated gene repression by controlling transcription initiation and burst frequency

**DOI:** 10.1101/2020.10.09.333294

**Authors:** Paula Dobrinić, Aleksander T. Szczurek, Robert J. Klose

## Abstract

The Polycomb repressive system plays a fundamental role in controlling gene expression during mammalian development. To achieve this, Polycomb repressive complexes 1 and 2 (PRC1 and PRC2) bind target genes and use histone modification-dependent feedback mechanisms to form Polycomb chromatin domains and repress transcription. The interrelatedness of PRC1 and PRC2 activity at these sites has made it difficult to discover the specific components of Polycomb chromatin domains that drive gene repression and to understand mechanistically how this is achieved. Here, by exploiting rapid degron-based approaches and time-resolved genomics we kinetically dissect Polycomb-mediated repression and discover that PRC1 functions independently of PRC2 to counteract RNA polymerase II binding and transcription initiation. Using single-cell gene expression analysis, we reveal that PRC1 acts uniformly within the cell population, and that repression is achieved by controlling transcriptional burst frequency. These important new discoveries provide a mechanistic and conceptual framework for Polycomb-dependent transcriptional control.

## Introduction

At the most basic level gene expression is controlled by DNA-binding transcription factors that recognise regulatory elements in the genome and shape how RNA polymerase functions at gene promoters. While this basic principle is shared across all clades of life, eukaryotes have evolved additional gene regulatory mechanisms that function through chromatin and often rely on post-translational modification of histone proteins ^1, 2^. However, the mechanisms through which these chromatin-based systems control transcription to enable gene regulation remains poorly understood and a major conceptual gap in our understanding of eukaryotic gene expression.

The importance of histone-modifying systems in gene regulation is exemplified by the Polycomb repressive system ^3, 4^, whose perturbation during early development causes catastrophic mis-regulation of gene expression and embryonic lethality ^5–8^. The Polycomb system is primarily composed of two large multi-protein complexes, Polycomb repressive complex 1 (PRC1) and 2 (PRC2). PRC1 is an E3 ubiquitin ligase that monoubiquitylates histone H2A at lysine 119 (H2AK119ub1) ^9–11^ and PRC2 is a histone methyltransfease that can mono-, di-, or trimethylate histone H3 at lysine 27 (H3K27me1/2/3) ^12–15^. PRCs function primarily at silent CpG island-associated gene promoters ^16–20^ where they form Polycomb chromatin domains that are characterised by H2AK119ub1, H3K27me3, and high-level occupancy of the Polycomb group proteins ^21, 22^. Importantly, the formation of Polycomb chromatin domains relies on feedback mechanisms between PRC1 and PRC2. For example, H2AK119ub1 placed by PRC1 is recognised by PRC2 ^23–25^ which deposits H3K27me3 that can in turn recruit more PRC1 ^12, 26, 27^. To initiate Polycomb chromatin domain formation, PRC1 and PRC2 utilise DNA-binding activities to engage with, or sample, promoter-associated CpG islands ^16–20^. These sampling activities identify lowly transcribed or inactive genes and enable Polycomb chromatin domain formation which then functions to counteract inappropriate gene expression and maintain normal cell identity ^3, 28^.

Despite an emerging view of how Polycomb chromatin domains form, we still have surprisingly little understanding of which components are required for gene repression and ultimately how the Polycomb system counteracts transcription. These fundamental questions have been difficult to answer due to the interrelated activities of PRC1 and PRC2 within Polycomb chromatin domains. For example, if one Polycomb complex is perturbed this inevitably disrupts the targeting and activity of the other, making it difficult to pinpoint the components of the system that are required for gene repression. Furthermore, genetic perturbation approaches currently used to disrupt Polycomb proteins are slow and asynchronous, making it almost impossible to study the primary effects that this system has on RNA polymerase II (Pol II) activity and transcription. Therefore, despite the Polycomb system being considered a paradigm for chromatin-based regulation of gene expression, we still have little mechanistic understanding of how it represses transcription to enable normal gene expression and development.

To uncover the component(s) of the Polycomb system necessary for gene repression and to discover how it counteracts Pol II-driven transcription, we have developed a degron system in mouse embryonic stem cells (ESCs) that leverages targeted proteolysis to rapidly and synchronously remove PRC1. By combining PRC1 depletion with time-resolved genomic analyses we discover that derepression of Polycomb target genes is remarkably quick and corresponds to active turnover of H2AK119ub1. Target gene derepression occurs in the presence of H3K27me3 and does not result from loss of PRC2 occupancy, demonstrating that PRC1 represses genes independently of PRC2.

Importantly, we discover that increases in Polycomb target gene transcription are dependent on new binding of Pol II and transcription initiation, not release of paused Pol II as has previously been proposed. Using single-cell gene expression analysis we reveal that PRC1 counteracts low-level gene expression uniformly throughout the cell population by controlling the frequency with which transcription bursts occur. Therefore, we discover a central role for PRC1 and H2AK119ub1 in controlling transcription initiation and bursting to define normal gene expression programmes and cell identity.

## Results

### Acute depletion of PRC1 reveals a rapid turnover of H2AK119ub1

Conditional knockout studies in mouse ESCs have shown that PRC1 is important for Polycomb chromatin domain formation and that its removal causes derepression of several thousand Polycomb target genes ^29–31^. However, in these systems loss of PRC1 and erosion of Polycomb chromatin domains is slow and asynchronous, making it difficult to pinpoint the components required for gene repression and to discover the primary mechanisms through which the Polycomb system inhibits transcription. To overcome these limitations, we have harnessed inducible proteolysis via the auxin-inducible degron (AID) system ^32, 33^ to rapidly deplete PRC1 through removing its core structural and catalytic subunit, RING1B (Figures 1A and S1A) ^34^. In this ESC system (PRC1^deg^) endogenously AID-tagged RING1B is expressed at wild type levels (Figure S1B) and PRC1 complexes form normally (Figure S1C). Moreover, RING1B is nearly completely degraded following 2 hours of auxin treatment (Figure 1B). In agreement with the requirement for RING1B in stabilising PRC1 complex formation ^29, 35, 36^, auxin treatment also caused a reduction in the levels of other PRC1 components (Figure S1D).

**Figure 1.**
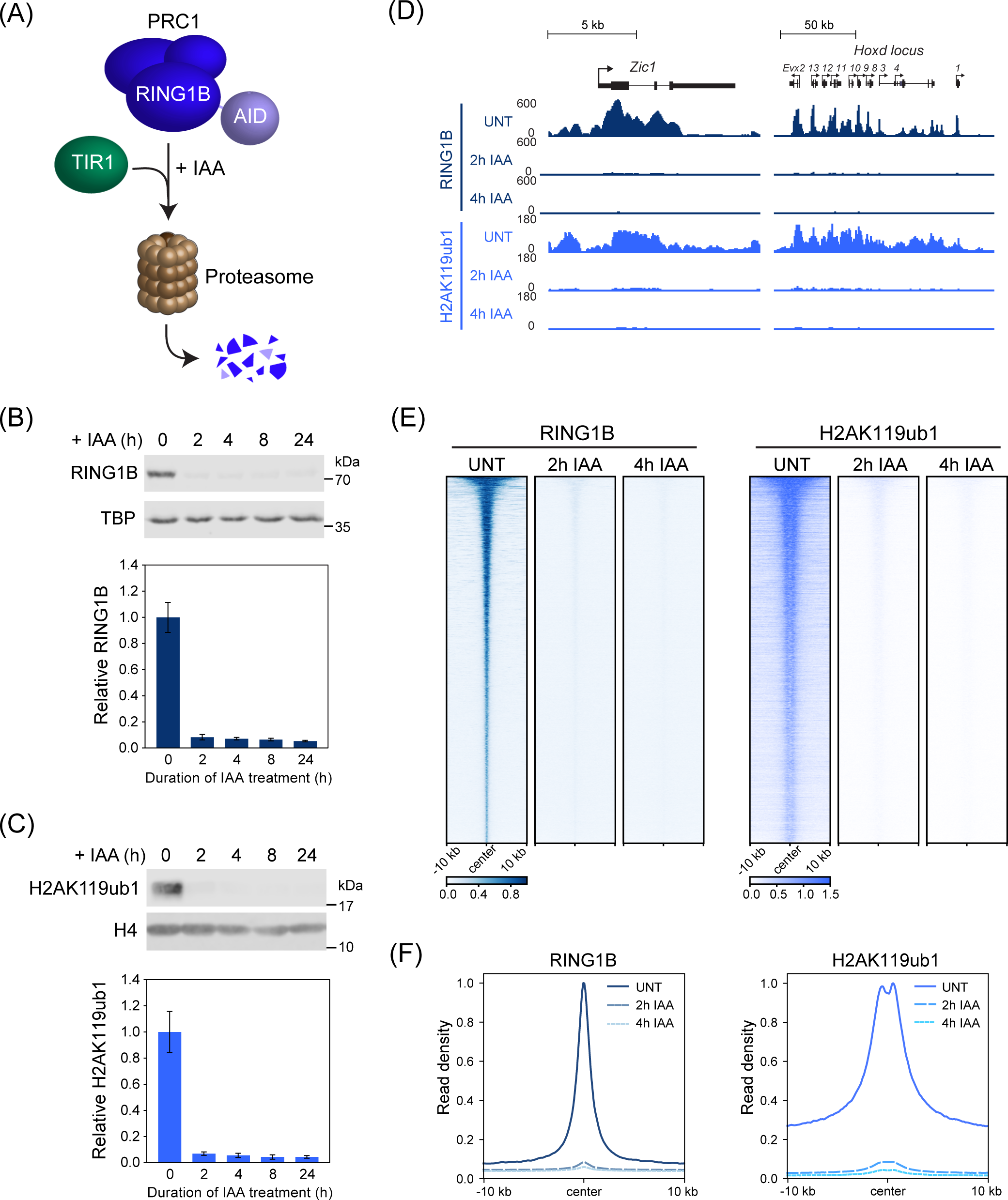
Acute depletion of PRC1 reveals a rapid turnover of H2AK119ub1. (A) A schematic illustrating the PRC1^deg^ system. Addition of auxin (IAA) induces proteasomal degradation of AID-RING1B. (B) Western blot analysis (upper panel) and quantification (lower panel) of RING1B in PRC1^deg^ cells treated with IAA for the indicated times. Data represents mean (n=2) ± SD. (C) Western blot analysis (upper panel) and quantification (lower panel) of H2AK119ub1 in PRC1^deg^ cells treated with IAA for the indicated times. Data represents mean (n=4) ± SD. (D) Genomic snapshots of typical Polycomb target genes, showing cChIP-seq signal for RING1B and H2AK119ub1 in PRC1^deg^ cells treated with IAA for the indicated times. (E) Heatmap analysis of RING1B (left) and H2AK119ub1 (right) cChIP-seq at RING1B-bound sites in PRC1^deg^ cells treated with IAA for the indicated times. Heatmaps were sorted by RING1B signal in untreated cells. (F) Metaplot analysis of data shown in (E).

Having demonstrated that we can rapidly remove PRC1, we first set out to determine how this affects H2AK119ub1, the histone modification placed by PRC1. Interestingly, western blot analysis revealed that H2AK119ub1 was barely detectable after only 2 hours of auxin treatment (Figure 1C) and calibrated chromatin immunoprecipitation coupled to massively parallel sequencing (cChIP-seq) demonstrated that RING1B and H2AK119ub1 were almost completely lost from PRC1 target sites and throughout the genome (Figures 1D, 1E, 1F and S1E). This indicates that H2AK119ub1 is rapidly turned over and that its deposition and removal are dynamically regulated. Importantly, the rapid kinetics of this system provides a new opportunity to dissect how PRC1 controls gene expression and discover which Polycomb chromatin domain components enable gene repression.

### Identification of primary PRC1 target genes reveals extremely rapid derepression following PRC1 removal

Our current understanding of PRC1-mediated gene repression has largely been derived from end point gene expression analyses following prolonged depletion of PRC1 using conditional knockout systems ^29–31^. As such, these experiments do not distinguish between primary and secondary gene expression effects, nor do they capture the kinetics of gene expression alterations as they occur at individual target genes. Therefore, using our rapid PRC1^deg^ system, we first wanted to identify the primary target genes repressed by PRC1. To achieve this, we carried out calibrated nuclear RNA-sequencing (cnRNA-seq) in a time course experiment following acute removal of PRC1. Remarkably, after only 2 hours of auxin treatment roughly one thousand (1044) genes increased in expression (Figure 2A). By 4 hours of auxin treatment this number had more than doubled (2841) and plateaued at 8 hours (4430). 94% of genes with increased expression at 2 hours of auxin treatment were Polycomb target genes and this class of genes also prominently featured at later time points (78% at 4 hours and 69% at 8 hours) (Figure 2B). Importantly, genes exhibiting increased expression at 2, 4, and 8 hours of auxin treatment showed a high degree of overlap (Figure 2C). This indicates that Polycomb target genes show different kinetics of derepression, with a subset of genes becoming derepressed very rapidly and relying heavily on PRC1 for their repression. Interestingly, after 24 hours of auxin treatment the number of derepressed genes was actually lower (2020) (Figure 2A), suggesting that gene expression adapts over time in response to the removal of PRC1, and reinforcing the importance of rapid depletion to identify the primary effects on gene expression. In agreement with this, the expression changes after 24 hours of auxin treatment correlated closely with the gene expression changes observed after long term conditional knockout of PRC1 (PRC1^cko^) ^30^ (Figure S2A). Therefore, our degron approach reveals primary Polycomb target genes and identifies a subset of genes that are particularly reliant on PRC1 to counteract their expression.

**Figure 2.**
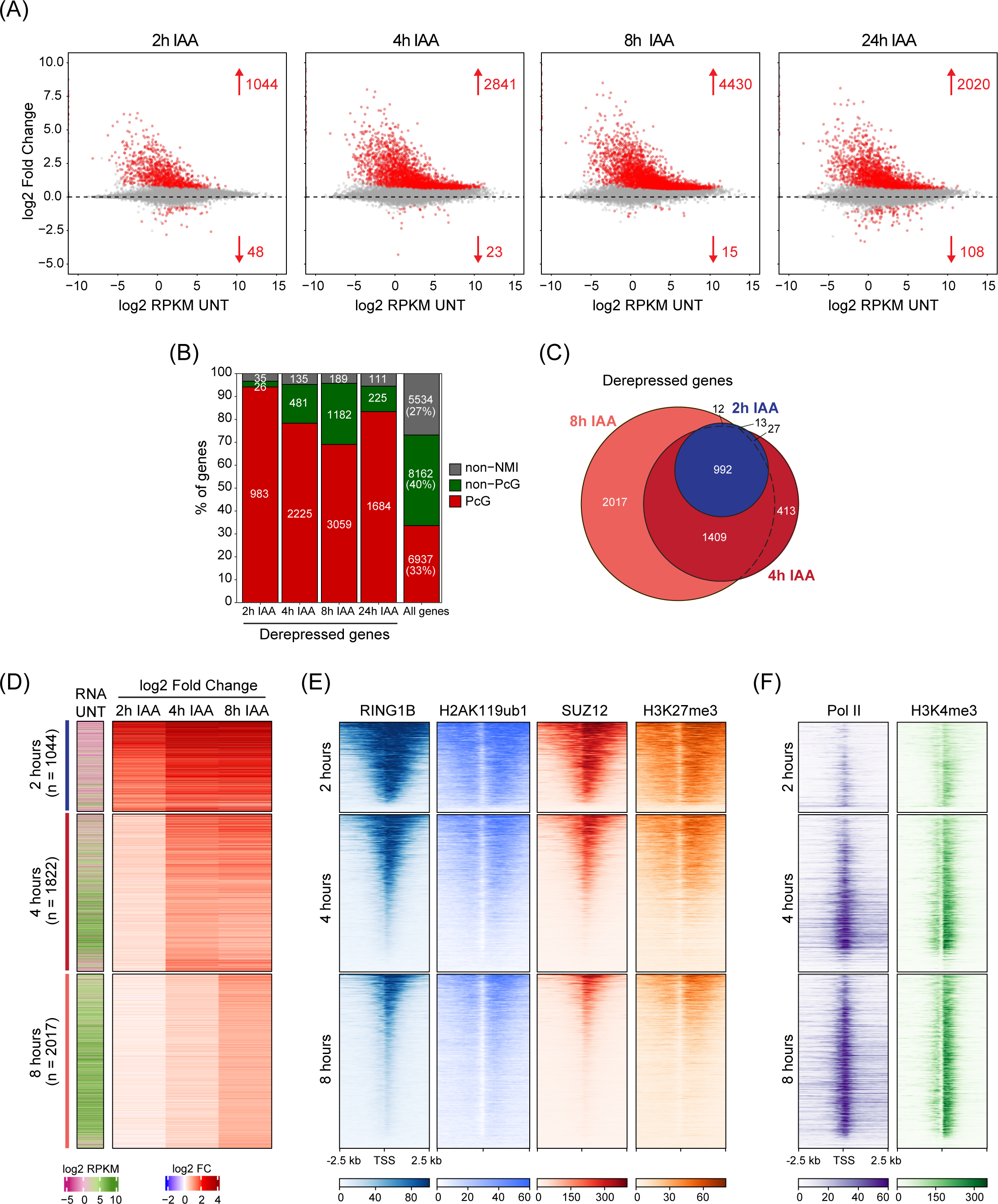
Perturbing PRC1 has immediate effects on gene expression. (A) MA plots depicting gene expression changes in PRC1^deg^ cells treated with IAA for the indicated times relative to untreated cells. Differentially expressed genes (padj < 0.05, fold change > 1.5) are labelled in red. (B) Distribution of different gene classes among genes with increased expression at different times following IAA treatment. Non-NMI, genes lacking a non-methylated CGI; Non-PcG, non-Polycomb-occupied genes; PcG, Polycomb-occupied genes. (C) A Venn diagram showing the overlap between significantly derepressed genes identified after treating cells with IAA for the indicated times. (D) Heatmaps depicting gene expression (cnRNA-seq) in untreated cells and changes following IAA treatment for the three groups of genes defined by the earliest time of derepression. Heatmaps are sorted by RING1B cChIP-seq signal as in (E). (E) Heatmaps showing cChIP-seq signal for Polycomb chromatin domain features (RING1B, H2AK119ub1, SUZ12, and H3K27me3) in untreated cells, at promoters of the gene groupings in (D). Heatmaps are sorted by RING1B cChIP-seq signal. (F) As in (E) but for total Pol II and H3K4me3 cChIP-seq.

To understand why some genes rely more heavily on PRC1 for their repression, we grouped genes based on the earliest time point they became derepressed (Figures 2D and S2B). Interestingly, among the three groups, genes affected by 2 hours had the lowest expression in untreated cells (Figures 2D and S2C), the largest magnitude of derepression after PRC1 removal, and tended to reach maximal expression by 4 hours of auxin treatment. In contrast, genes that displayed significant increases in expression at either 4 or 8 hours showed higher initial expression and a substantially smaller magnitude of derepression. When we explored the chromatin features associated with the promoters of these different groups of genes in untreated cells, those that became derepressed at 2 hours had the highest levels of Polycomb chromatin domain features including RING1B, H2AK119ub1, SUZ12, and H3K27me3 (Figures 2E and S2D). They also displayed the lowest levels of Pol II and transcription-associated histone modification H3K4me3 (Figures 2F and S2E). Conversely, genes derepressed at 4 and 8 hours tended to have lower levels of Polycomb chromatin domain features and much higher Pol II and H3K4me3, indicating that these genes are already partially transcribed. Therefore, our new time-resolved gene expression analysis reveals that PRC1 plays a central role in repressing Polycomb target genes with low occupancy of the transcriptional machinery, and that these genes become rapidly derepressed in the absence of PRC1.

### PRC1-mediated gene repression does not rely directly on PRC2 or H3K27me3

The lack of synchrony and temporal resolution of classical genetic perturbation approaches coupled with the interrelatedness of PRC1 and PRC2 function at Polycomb target genes has made it almost impossible to pinpoint the components of Polycomb chromatin domains that are required for gene repression. Having established that H2AK119ub1 is lost very rapidly following PRC1 removal, we set out to determine whether the rapidity of this turnover would provide a new window of opportunity to dissect the relationship between PRC1 function, Polycomb chromatin domain features, and gene repression. To achieve this, we first carried out time-resolved cChIP-seq analysis for SUZ12, a core structural component of PRC2, following PRC1 depletion. Remarkably, this revealed that PRC2 binding was majorly reduced after only 2 hours of auxin treatment at promoters of derepressed genes (Figures3A, 3B and S3A). Importantly, these reductions in PRC2 binding occurred in concert with loss of H2AK119ub1 and changed only modestly thereafter, consistent with PRC2 relying on binding to H2AK119ub1 for occupancy at Polycomb chromatin domains ^23–25, 37^. However, previous studies have also indicated that displacement of PRC2 from chromatin can be caused by gene transcription through PRC2 binding to nascent RNA ^38–40^. To ensure that the effects we observed on PRC2 occupancy were not solely due to alterations in gene expression, we compared PRC2 binding at Polycomb target genes that were either unaffected or increased in expression after 2 hours of auxin treatment. Importantly, major reductions in SUZ12 binding occurred regardless of whether the associated gene displayed changes in expression (Figure S3B). Together this reveals that PRC2 binding to chromatin is highly dependent on PRC1/H2AK119ub1.

Based on the rapid reductions in PRC2 binding we observed in the absence of PRC1 and H2AK119ub1, we were curious to examine how this affected H3K27me3. Therefore, we carried out cChIP-seq for H3K27me3 over the same time course. Despite major reductions in PRC2 binding, there was a much more modest effect on H3K27me3 levels at 2 hours of auxin treatment and over the remainder of the time course we observed a slow reduction in H3K27me3 levels (Figures 3A, 3C and S3C). The rate at which H3K27me3 reductions occurred was consistent with an exponential decay model based on dilution during replication and cell division, as opposed to active turnover (Figure S3D). Given that PRC2 protein levels remained unaffected by PRC1 removal (Figure S1D), PRC2-H3K27me3 read-write mechanisms ^41–43^ alone must be insufficient to reinstate H3K27me3 at Polycomb chromatin domains following DNA replication ^44^. Instead, stabilisation of PRC2 at Polycomb chromatin domains via binding to H2AK119ub1 must contribute centrally to epigenetic maintenance of H3K27me3 during DNA replication.

**Figure 3.**
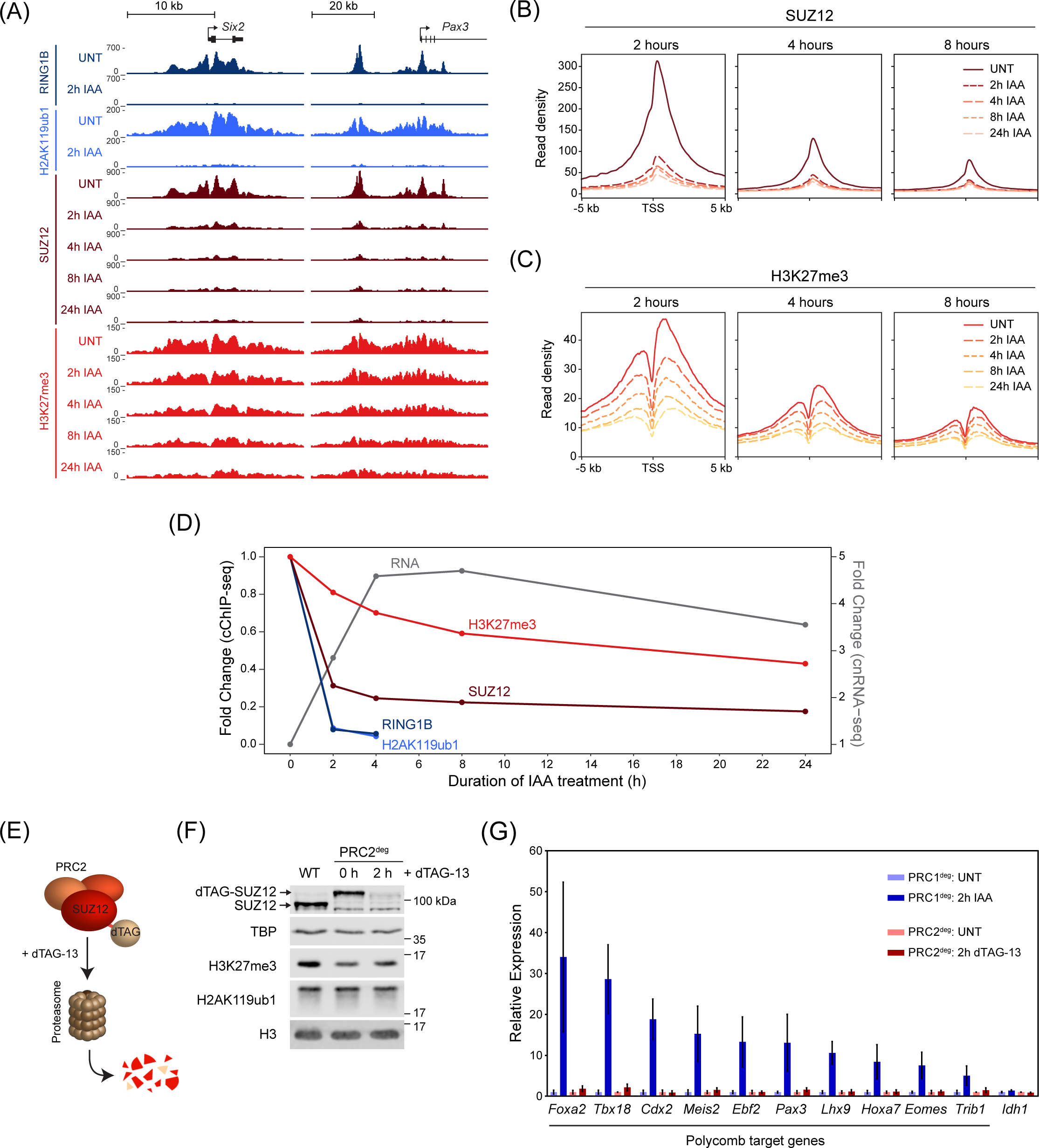
PRC1-mediated gene repression does not rely directly on PRC2 or H3K27me3. (A) Genomic snapshots of typical Polycomb target genes, showing cChIP-seq signal for RING1B, H2AK119ub1, SUZ12 and H3K27me3 in PRC1^deg^ cells treated with IAA for the indicated times. (B) Metaplot analysis of SUZ12 cChIP-seq at promoters of the three groups of genes defined by the earliest time of derepression in PRC1^deg^ cells treated with IAA for the indicated times. (C) As in (B) for H3K27me3 cChIP-seq. (D) Dynamics of gene expression changes in comparison with the reduction in cChIP-seq signal (TSS ± 2.5 kb) for Polycomb factors and associated histone modifications. Shown are median changes relative to untreated cells for genes derepressed after 2h of IAA treatment (n = 1044). (E) Schematic illustration of the PRC2^deg^ system. Endogenous SUZ12 is N-terminally tagged and degraded by the proteasome after addition of the dTAG-13 compound. (F) Western blot analysis of SUZ12, H3K27me3 and H2AK119ub1 in wild type (WT) and PRC2^deg^ cells (untreated and treated with dTAG-13 compound). (G) RT-qPCR analysis of expression for a panel of Polycomb target genes in PRC1^deg^ and PRC2^deg^ cells treated for 2 hours with IAA or dTAG-13, respectively. Gene expression is normalised to *U6 snRNA* and shown relative to the average expression in untreated cells for each individual cell line. Data represents mean (n=3) ± SEM.

Interestingly, these analyses also revealed that derepression of Polycomb target genes at 2 hours of auxin treatment occurred despite the fact that H3K27me3 is still largely retained. This suggests that gene expression can occur despite the presence of this modification and that PRC2 catalytic activity might therefore contribute little to the repressive nature of Polycomb chromatin domains (Figure 3D). However, at this time point PRC2 occupancy was also majorly reduced, making it difficult to exclude a contribution of PRC2 binding to gene repression. To examine this possibility, we utilised a PRC2 degron system (PRC2^deg^) where treatment with the dTAG-13 compound for 2 hours results in the near complete removal of SUZ12, without affecting H3K27me3 or H2AK119ub1 (Figure 3E, F)^45, 46^. We then compared changes in expression of Polycomb target genes in the PRC2^deg^ and PRC1^deg^ cells. While removal of PRC1 caused derepression of Polycomb target genes, removal of PRC2 had no effect (Figure 3G)^47^, demonstrating that reductions in PRC2 occupancy following PRC1 removal do not account for the increased expression of Polycomb target genes. Together, our rapid loss-of-function systems and time-resolved analysis reveal that PRC2 occupancy at Polycomb chromatin domains is primarily defined by PRC1/H2AK119ub1, and that PRC1-mediated gene repression does not rely directly on PRC2 or H3K27me3.

### Removal of PRC1 causes a rapid new binding of Pol II

Despite the essential roles that the Polycomb system plays in regulating gene expression, our understanding of how it counteracts transcription to achieve repression has remained enigmatic. In fact, previous work has implicated the Polycomb system in regulating various and distinct phases of transcription, including initiation, pausing/priming, and elongation ^48–54^. Having shown that we can identify the earliest and most primary effects on gene expression following removal of PRC1, we set out to discover how the Polycomb system counteracts the process of transcription at its target genes. To begin addressing this question, we used an antibody that recognises total Pol II and carried out cChIP-seq in a time course experiment following removal of PRC1. This revealed that there was a rapid new binding of Pol II at Polycomb target gene promoters, and this was most pronounced at genes that show increased expression at 2 hours after removal of PRC1 (Figures 4A, 4B and S4A). Remarkably, despite the fact that expression continued to increase for many of these genes throughout the time course, maximum levels of new Pol II binding had already occurred at 2 hours (Figures 4B and S4B). In fact, even at genes where significant changes in expression did not manifest until 4 or 8 hours after auxin treatment maximal new promoter-proximal binding of Pol II was evident at 2 hours. This indicates that the underlying defect in Polycomb chromatin domain function that enables new Pol II binding had occurred by 2 hours of auxin treatment, and that the observed delay in measuring significant gene expression changes at these genes is likely due to the time it takes for multiple rounds of transcription to occur and new transcripts to accumulate. In agreement with this possibility when we examined H3K4me3, a histone modification that is associated with active transcription ^55^, it continued to accumulate until 4 hours after auxin treatment (Figures 4A and 4C), regardless of the time at which genes became significantly derepressed (Figures S4B and S4C). In conclusion, new binding of Pol II closely follows PRC1/H2AK119ub1 removal, with H3K4me3 accumulating more slowly as transcription proceeds (Figure 4D). Together these observations indicate that PRC1 functions to counteract Pol II binding at Polycomb target gene promoters.

**Figure 4.**
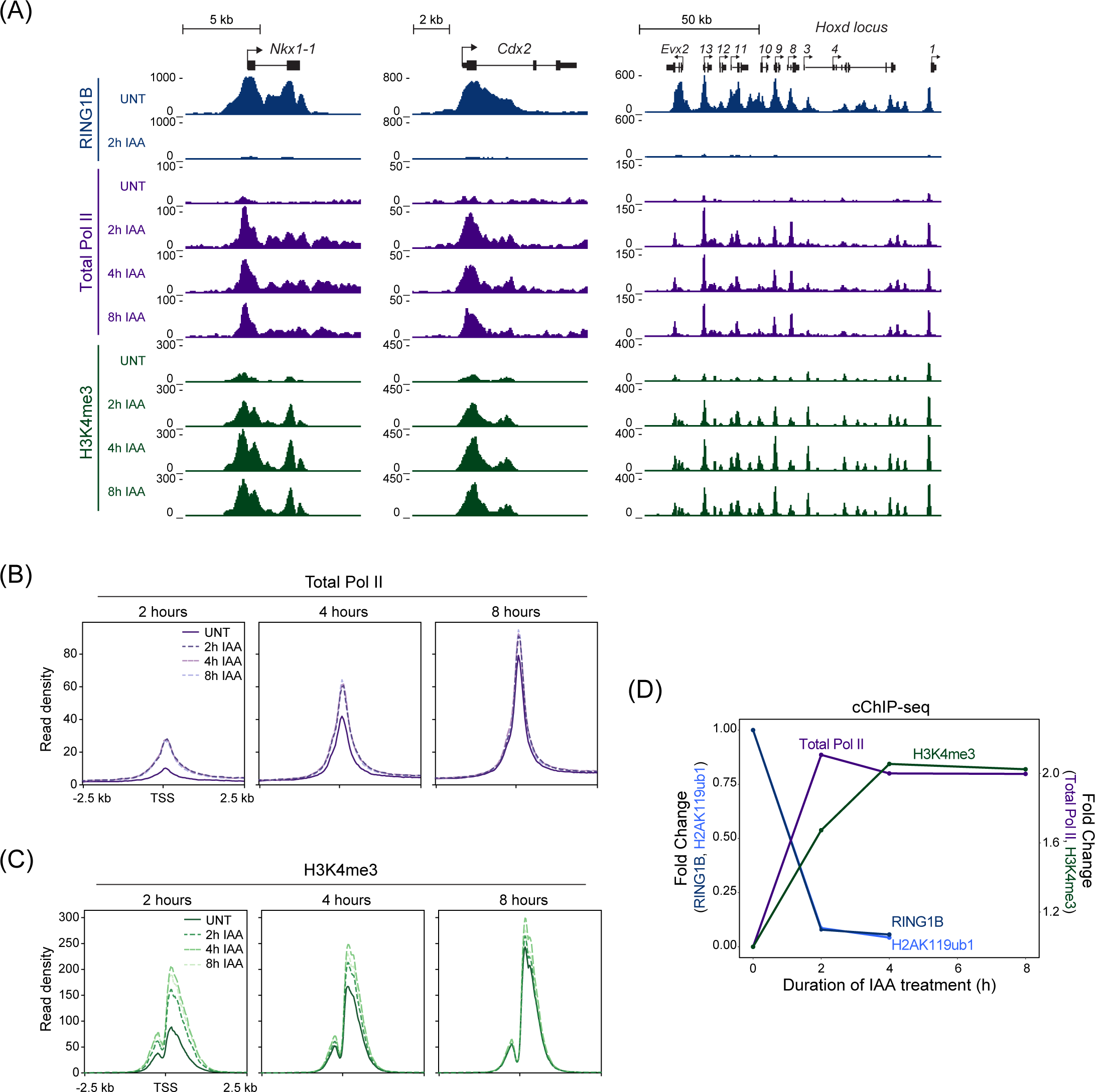
Removal of PRC1 causes a rapid new binding of Pol II and accumulation of H3K4me3. (A) Genomic snapshots of typical Polycomb target genes, showing cChIP-seq signal for RING1B, total Pol II and H3K4me3 in PRC1^deg^ cells treated with IAA for the indicated times. (B) Metaplot analysis of total Pol II cChIP-seq at promoters of the three groups of genes defined by the earliest time of derepression in PRC1^deg^ cells treated with IAA for the indicated times. The profiles represent the median signal over the shown genomic region. (C) As in (B) but for H3K4me3 cChIP-seq. (D) Dynamics of decreases in cChIP-seq signal for RING1B and H2AK119ub1 and increases in total Pol II and H3K4me3 signal at promoters (TSS ± 2.5 kb) of genes derepressed after 2h of IAA treatment (n = 1044). Shown are median changes relative to untreated cells.

### New initiation is required for Polycomb target gene expression

The phosphorylation state of the C-terminal heptapeptide repeat domain (CTD) of Pol II is often used to characterise distinct stages of transcription. For example, phosphorylation of serine 5 (Ser5P) on the CTD is usually associated with initiated or promoter-proximal paused Pol II and serine 2 phosphorylation (Ser2P) is associated with transcription elongation ^56^. Previous studies examining Pol II phosphorylation states in ESCs have reported high levels of Ser5P-Pol II at Polycomb target gene promoters ^50, 57^ despite their lowly, or non-transcribed, state. Based on these observations it was proposed that the Polycomb system functions to repress gene expression by keeping Pol II in a poised state, and that release of poised Pol II in the absence of PRC1 was responsible for Polycomb target gene derepression ^50, 57^. However, subsequent analyses have suggested that Polycomb target genes have extremely low levels of transcriptionally engaged Pol II at their promoters ^58, 59^ and we found they were characterised by low levels of total Pol II binding in our ChIP-seq analysis. These findings are therefore inconsistent with the release of poised Pol II being sufficient to cause derepression of Polycomb target genes in the absence of PRC1.

Given our capacity to rapidly remove PRC1 and capture the earliest events leading to gene derepression, we set out to examine in more detail Ser5P-Pol II at Polycomb target genes and to understand how it is related to their derepression. To achieve this, we carried out cChIP-seq for Ser5-Pol II in untreated cells and after removal of PRC1. As described previously, Ser5P-Pol II ChIP-seq signal was present at Polycomb target genes in untreated cells ^50, 57^, despite extremely low levels of total Pol II signal (Figures 5A and S5A). However, compared to actively transcribed genes where Ser5P-Pol II occurs just downstream of the transcription start site, the distribution of Ser5P-Pol II at Polycomb target genes was highly atypical and mirrored the distribution of the underlying Polycomb chromatin domain, which in some instances extended tens of kilobases outside of the gene itself (Figure 5A) ^50, 57^.

**Figure 5.**
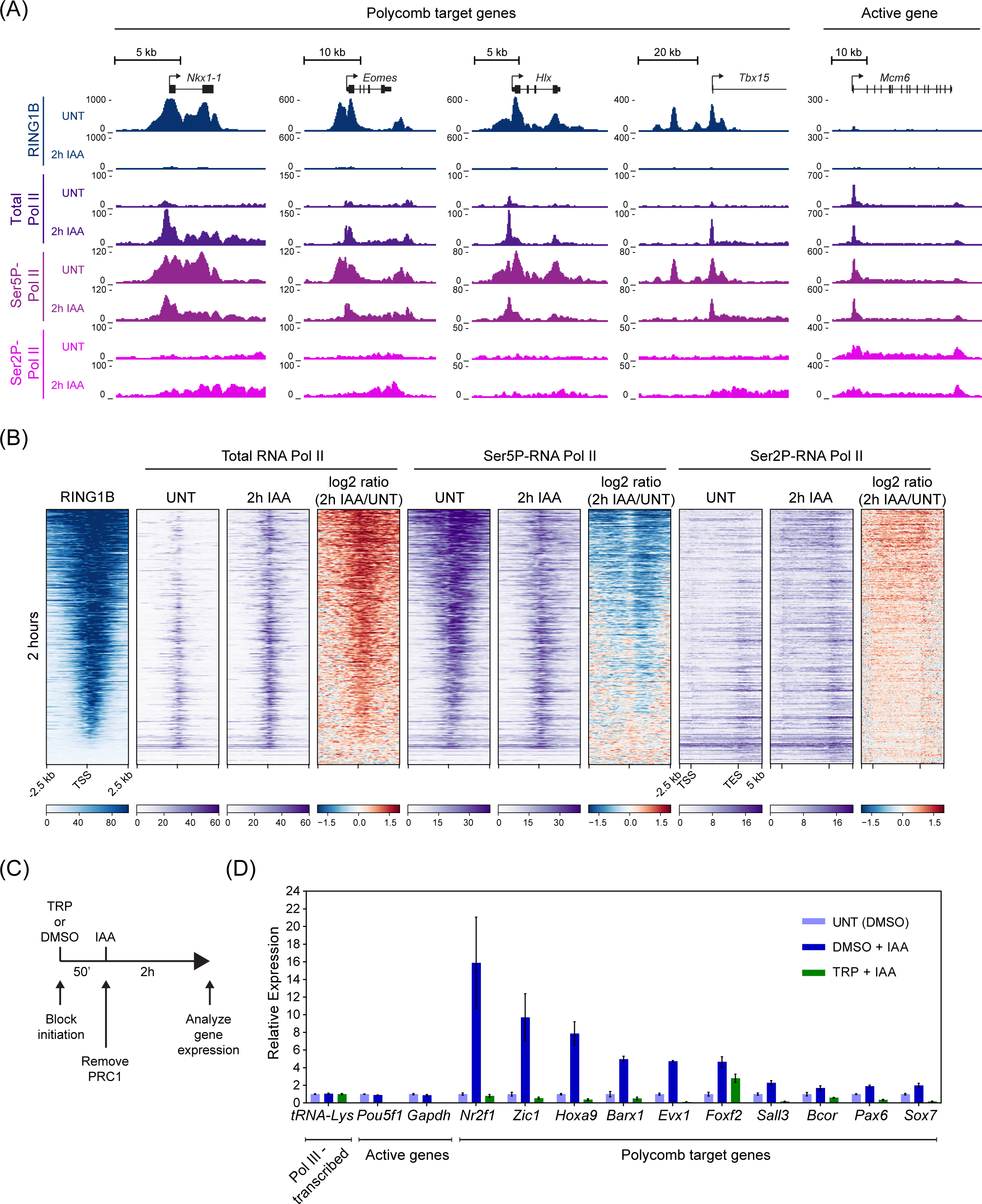
New initiation is required for Polycomb target gene derepression. (A) Genomic snapshots of typical Polycomb target loci and one highly transcribed gene (*Mcm6*), showing cChIP-seq signal for RING1B, total Pol II, Ser5P-Pol II and Ser2P-Pol II in untreated PRC1^deg^ cells and 2 hours after IAA addition. (B) Heatmaps illustrating RING1B, total Pol II, Ser5P-Pol II and Ser2P-Pol II levels and changes in polymerase occupancy following 2h of IAA treatment for genes derepressed at that point (n = 1044). Heatmaps are sorted by RING1B signal in untreated cells. All derepressed Polycomb targets are shown in Figure S5B. (C) Schematic of the experimental setup to test dependency of Polycomb target gene derepression on new initiation. 50 min prior to PRC1 depletion, cells are pre-treated with triptolide (TRP), an inhibitor of Pol II initiation, or DMSO. PRC1 removal is induced by IAA for 2 hours before collecting cells for the analysis. (D) RT-qPCR analysis of gene expression changes in PRC1^deg^ cells following 2h of IAA treatment, with or without 50 min pre-treatment with TRP. Gene expression is normalised to *U6 snRNA* and shown relative to the average expression in untreated cells. Data represents mean (n=3) ± SEM.

This was most evident when we focussed on genes that are derepressed at 2 hours and have large Polycomb chromatin domains (Figure 5B). Interestingly, after PRC1 removal, this atypical Ser5P-Pol II distribution was lost and now Ser5P-Pol II signal was focused at transcription start sites corresponding to where new total Pol II binding was observed, presumably reflecting new transcription initiation (Figures 5A, 5B, S5A, S5B and S5C). We also observed small increases in Ser2P-Pol II levels in the body of derepressed genes, in agreement with their increased transcription (Figures 5A, 5B, S5A, S5B and S5D). Together these observations are consistent with rapid Polycomb target gene derepression after PRC1 removal relying on new Pol II binding and transcription initiation, not release of a paused/poised polymerase. To test this more directly, we treated cells with triptolide, a small molecule inhibitor that disrupts initiation by Pol II, removed PRC1, and then asked if rapid derepression of Polycomb target genes could occur in the absence of new transcription initiation (Figure 5C). As expected, triptolide treatment did not affect the transcription of a RNA polymerase III (Pol III)-transcribed gene, but efficiently blocked Pol II-mediated transcription of active genes. More importantly, derepression of almost all analysed Polycomb target genes was abolished in triptolide-treated cells (Figure 5D). Therefore, we conclude that new transcription initiation, not pause release, explains the rapid derepression of Polycomb target genes in the absence of PRC1 and demonstrates that PRC1 primarily functions to counteract transcription initiation.

### The Polycomb system constrains low-level gene expression but does not block its target genes from switching into a highly expressed or activated state

Our genomic experiments suggest that Polycomb system functions through PRC1 and H2AK119ub1 to counteract the process of transcription initiation. However, the ensemble nature of genomic experiments makes it impossible to understand if these effects occur in all cells in the population, or just in a subset of cells. Furthermore, they lack the resolution to define what aspect of transcription initiation is controlled by the Polycomb system. Therefore, to discover how the Polycomb system regulates gene expression in single cells, we employed single-molecule RNA fluorescent in situ hybridization (smRNA-FISH) (Figure 6A) and developed a high-throughput imaging and analysis approach to enable transcript counting (Figure S6). We then used this approach to study expression levels of 16 Polycomb target genes in single cells before and after PRC1 removal (Figures 6B, S7A and S7B). Interestingly, we found that all of the Polycomb target genes analysed were expressed to some degree, but most had very low transcript numbers (∼1 transcript/cell) (Figure 6B). A subset (*E2f6*, *Gbx2,* and *Zic2*) displayed higher average transcript numbers (∼5 to ∼12 transcripts/cell) but these were still much lower than actively transcribed control genes (∼33 and ∼38 transcripts/cell for *Hspg2* and *Tfrc*, respectively) (Figure 6B). Therefore, Polycomb target genes are expressed, albeit very lowly, demonstrating that the Polycomb system must not be an impervious block to gene transcription.

**Figure 6.**
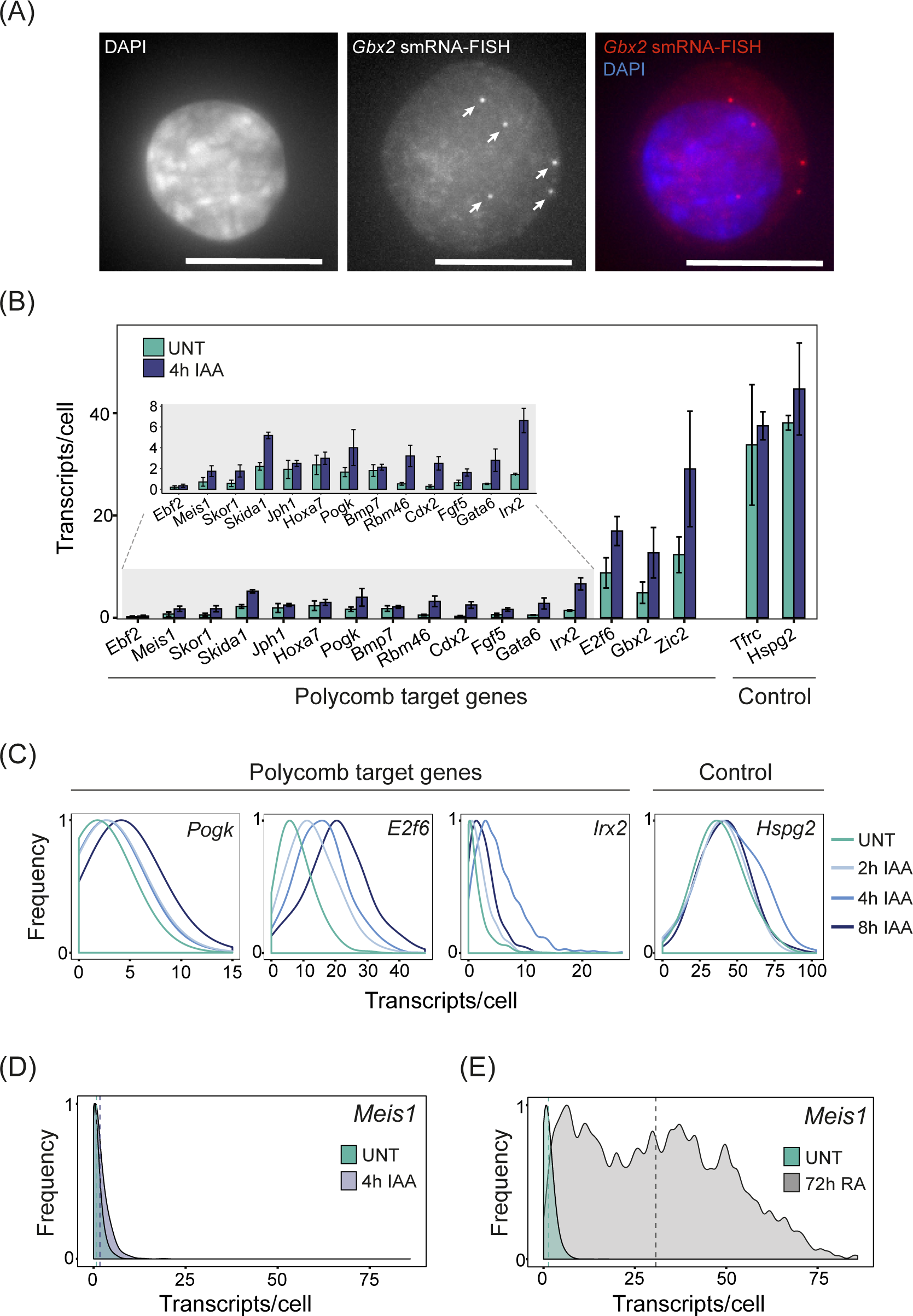
Single-cell analysis of Polycomb target gene expression. (A) A smRNA-FISH image of the Polycomb target gene *Gbx2*. Individual dots indicated with arrows correspond to single transcripts of *Gbx2*. DAPI is shown to indicate the nucleus (left panel). Maximal projections of 3D stacks are presented and the scale bar corresponds to 10 µm. (B) A bar graph illustrating the mean number of transcripts per cell in PRC1^deg^ cells before and after 4 hours of IAA treatment to remove PRC1 for 16 Polycomb target genes and two control genes. Data represents mean (n=3) ± SD. (C) Normalised density plots illustrating the distribution of transcripts across the cell population in PRC1^deg^ cells treated with IAA for the indicated times for selected Polycomb target genes and a control gene. All analysed genes are shown in Figure S7C. (D) As in (C) but for *Meis1* in untreated PRC1^deg^ cells and after 4 hours of IAA treatment to remove PRC1. The vertical dashed lines correspond to the mean value of transcripts per cell for the respective condition. (E) As in (D) but for *Meis1* in untreated ESCs and after 72 hours of retinoic acid treatment (RA).

Furthermore, this indicates that Polycomb target genes must be subject to low-level transcriptional signals that can lead to some level of gene expression.

We were then curious to examine how transcript numbers changed following removal of PRC1. Importantly, this revealed that Polycomb target genes now transitioned into a more expressed state (Figure 6B). However, given their very low initial expression levels, even genes that showed a reasonably large increase in expression after 4 hours of auxin treatment (e.g. *Irx2*, ∼5-fold) still only accumulated on average ∼7 transcripts per cell, far below the levels that we observed for control genes (Figure 6B). This modest effect on absolute transcript number was evident for most genes across the time course of PRC1 removal (Figures S7D and S7E). Furthermore, when we asked how the effects on transcript levels in individual cells accounted for increases in Polycomb target gene expression, it was clear that almost all cells in the population displayed small increases in expression, as opposed to only a subset of cells switching into a highly expressed or activated state (Figures 6C and S7C). Therefore, PRC1 primarily counteracts low-level transcriptional signals that are present across the cell population, as opposed to blocking strong transcription signals that could activate gene expression either in single cells or more broadly.

Interestingly, in contrast to what we observe here, previous single-cell RNA-sequencing (scRNA-seq) analysis had concluded that Polycomb target genes, and particularly those that were more highly expressed, show a bimodal distribution of transcript levels in the cell population ^60^. It was proposed that this bimodality arose from some cells displaying very low Polycomb target gene expression and others existing in a highly expressed state. Furthermore, it was reported that increases in gene expression after removal of PRC1 resulted from lowly expressing cells switching into a highly expressed or activated state ^60^. Importantly, we did not observe this type of bimodality for Polycomb target genes in untreated cells, nor did it emerge following removal of PRC1 (Figures 6C and S7C). This difference is likely explained by scRNA-seq lacking the capacity to accurately quantify lowly expressed genes ^61^ and absolute transcript numbers. These limitations are overcome by our smRNA-FISH which enables direct quantification with single-transcript sensitivity. However, to further validate that the effects on gene expression that we observed in single cells following removal of PRC1 did not result from lowly expressing cells switching into a highly expressed or activated state, we focussed on the *Meis1* gene. When PRC1 was removed, *Meis1* expression increased by 2.5-fold (0.7 to 1.75 transcripts/cell) (Figure 6D). In contrast, when cells were treated with retinoic acid to induce differentiation, *Meis1* expression increased more than 40-fold and switched into a highly expressed state with transcript numbers similar to other actively transcribed genes (∼30 transcripts/cell) (Figure 6E). This observation reinforces our conclusion that the Polycomb system primarily functions to counteract low-level transcriptional signals present in most cells in the population, as opposed to protecting its target genes from switching into a highly expressed or activated state.

### The Polycomb system represses gene expression by limiting transcription burst frequency

How the Polycomb system and other chromatin-based repressors function to regulate the process of transcription at gene promoters remains very poorly understood and constitutes a major conceptual gap in the field. Our single-cell analysis revealed that Polycomb target genes are expressed, albeit at low levels, and that PRC1 functions to counteract this expression in most cells in the population. This indicates that Polycomb target genes are constantly subject to signals that promote transcription and therefore we hypothesized that PRC1 must function to counteract some fundamental aspect of the transcription process at these gene promoters. In mammals the process of transcription from gene promoters is pulsatile and stochastic ^62, 63^. During periods of activity, individual genes undergo bursts of transcription initiation and elongation, leading to production of multiple transcripts. This is often followed by prolonged periods of inactivity. The frequency with which transcription bursts occur (burst frequency) and how many transcripts are produced during a burst (burst size) underpin gene expression (Figure 7A).

**Figure 7.**
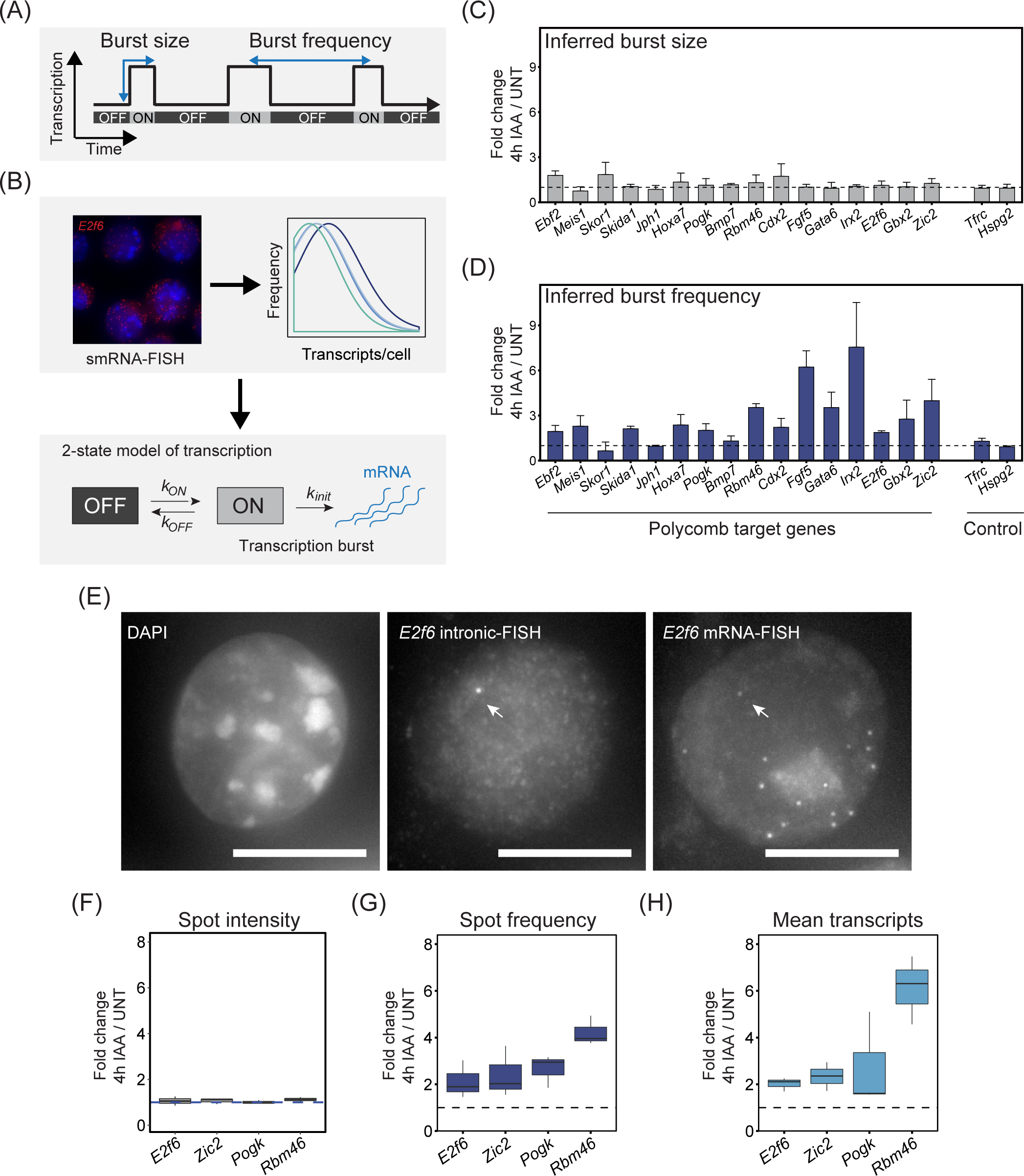
The Polycomb system represses gene expression by limiting transcription burst frequency. (A) A schematic illustrating the stochastic and pulsatile nature of gene transcription. Over time, a gene promoter can exist in an OFF state where no transcription is occurring or ON state where multiple transcripts are produced (a burst of transcription). The number of transcripts that are produced in a burst is referred to as the burst size and the burst frequency is the time between transcription bursts. Importantly, transcription burst sizes and their frequency are central determinants of gene expression. (B) A schematic illustrating how population level transcript distributions from smRNA-FISH are used to inform a two state model of transcription and infer kinetic parameters of transcription. (C) A bar graph illustrating the fold change in inferred transcription burst size for Polycomb target and control genes. Data represents mean (n=3) ± SD. The dashed horizontal line corresponds to no change. (D) As in (C) but for inferred burst frequency. (E) An example of a nascent smRNA-FISH (intronic FISH) image for *E2f6*, with an individual bright dot corresponding to a nascent transcription spot indicated with an arrow (middle panel). Standard smRNA-FISH (targeting exonic sequences) in the same cells illustrates individual transcripts and the nascent transcription spot (right panel). DAPI is shown to indicate the nucleus (left panel). The scale bar corresponds to 10 µm. (F) Boxplots illustrating the fold change in nascent transcription spot intensity for the indicated Polycomb target genes following 4h IAA treatment to remove PRC1. The dashed horizontal line corresponds to no change. (G) As in (F) but fold change in nascent transcription spot frequency. (H) As in (F) but fold change in the mean number of transcripts per cell.

Therefore, we set out to characterise these features of Polycomb target gene transcription and discover how they are controlled by PRC1 to enable gene repression. To achieve this, we used transcript distributions from our smRNA-FISH measurements and a 2-state model of transcription to infer the kinetic parameters of Polycomb target gene transcription ^62, 64–66^ (Figures 7B, S8A and S8B). In untreated cells the majority of Polycomb target genes had very small transcription burst sizes (∼1-3 transcripts) compared to highly expressed control genes (∼7-15 transcripts) (Figure S8C). However, some Polycomb target genes with higher expression levels (*Gbx2* and *Zic2)* in untreated cells also displayed larger burst sizes (∼6-17 transcripts) (Figure S8C). To determine if PRC1 controls this aspect of transcription we examined how burst size is affected when PRC1 is removed. Importantly, despite some genes increasing in expression by up to 10-fold (Figures 6B and S7D), the burst size of most Polycomb target genes was unaffected by PRC1 removal (Figure 7C). In instances where there were very modest effects on burst size (*Ebf2, Skor1, Cdx2*) (Figure S8E), these alone were insufficient to explain the corresponding increase in transcripts (Figure S7D). Together this suggests that PRC1 does not primarily function to regulate transcription burst size.

Having established that PRC1 removal has little effect on transcription burst size, we reasoned that transcriptional repression by PRC1 may function through limiting burst frequency. To test this, we revisited the two-state model of our smRNA-FISH measurements to estimate transcription burst initiation rate (kON), which can be used as a proxy for burst frequency. This revealed that the vast majority of Polycomb target genes displayed increases in burst frequency when PRC1 was removed (Figure 7D) and this was in line with the increased expression of these genes (Figures 6B and S7D). Therefore, our modelling-based approaches suggest that PRC1 primarily limits burst frequency to repress gene expression.

To test this possibility more directly, we used intronic smRNA-FISH that captures short-lived nascent pre-mRNAs as they accumulate at the site of active transcription and therefore more directly interrogates promoter activity (Figure 7E). The number of nascent transcription spots in the cell population represents a measure of burst frequency, while the intensity of individual spots provides information about transcription burst size ^67^. We designed intronic FISH probes for four Polycomb target genes, two of which were lowly expressed (*Rbm46* and *Pogk*), and two more highly expressed (*E2f6* and *Zic2*). When we measured spot intensities for these genes in untreated cells they scaled well with inferred burst sizes (Figure S8D) and, importantly, they remained unchanged following removal of PRC1, in line with our conclusions from modelling of smRNA-FISH transcript distributions (Figures 7F and S8E). However, when we examined the number of nascent transcription spots in the cell population after PRC1 removal, we observed clear increases in their frequency for all four genes (Figure 7G) and this effect corresponded well to the increases in expression (Figure 7H). Therefore, our new single-cell measurements reveal that PRC1 predominantly represses gene expression by limiting transcription burst frequency, not burst size, and uncovers how the Polycomb system affects transcription to regulate gene expression.

## Discussion

Despite decades of intense study in many different contexts ^4^, at least two fundamental questions about how the Polycomb system functions in gene regulation have remained elusive, namely, which component(s) of Polycomb chromatin domains are required for gene repression and how do these features regulate transcription to elicit gene repression. Here, using ESCs as a model system and leveraging rapid depletion approaches coupled with time-resolved calibrated genomics and single-cell gene expression analysis we address these critical questions. Importantly, we reveal that acute removal of PRC1 results in a rapid loss of H2AK119ub1 and derepression of Polycomb target genes (Figure 1 and 2). This occurs independently of PRC2 and H3K27me3 (Figure 3), demonstrating that Polycomb chromatin domain-mediated repression primarily relies on PRC1. We discover that PRC1-dependent gene repression functions by controlling Pol II binding and transcription initiation from gene promoters (Figure 4), not pause release as previously proposed (Figure 5). Single-cell analysis reveals that this repressive activity functions uniformly across the cell population (Figure 6) to limit the frequency of transcriptional bursts from Polycomb target genes (Figure 7). Together these new discoveries demonstrate that the Polycomb system uses PRC1 and H2AK119ub1 to control PoI II binding and limit the frequency of transcription initiation events to repress its target genes.

The interrelatedness of Polycomb repressive complex activities at target gene promoters has made identifying the mechanisms of gene repression extremely challenging. Using rapid depletion and time-resolved genomic analyses we now show that Polycomb target genes become immediately derepressed when PRC1 and H2AK119ub1 are lost from Polycomb chromatin domains, and that derepression occurs despite the presence of H3K27me3. Furthermore, rapid removal of PRC2 has no effect on Polycomb target gene repression. Therefore, these experiments narrow down the repressive activity of Polycomb chromatin domains to PRC1, but alone they do not define whether PRC1 complexes or H2AK119ub1 are responsible for transcriptional repression. Previously we and others developed ESCs systems where we could inducibly inactivate PRC1 catalysis, but leave PRC1 complexes intact and still able to bind target genes ^29, 68^. These experiments showed that catalysis by PRC1 is essential for gene repression, but these effects were confounded by the fact that H3K27me3 and PRC2 occupancy were also profoundly affected due to the aforementioned feedback mechanisms and the protracted time required to convert PRC1 into a catalytically deficient form. Our rapid perturbation experiments now enable us to exclude a central and direct role for PRC2 and H3K27me3 in PRC1-dependent gene repression, therefore strongly implicating H2AK119ub1 as the defining feature of Polycomb chromatin domains that enables gene repression in ESCs. These observations are in agreement with previous findings that canonical PRC1 complexes that bind to chromatin through H3K27me3, but which have limited capacity to deposit H2AK119ub1 ^23, 69–71^, contribute little to gene repression ^72–76^. In contrast, variant PRC1 complexes that deposit almost all H2AK119ub1 are essential for gene repression ^30^. Furthermore, when H2AK119ub1 is directly incorporated into chromatinised gene promoter templates *in vitro,* this specifically inhibits transcription ^49, 77^. Together these observations suggest that H2AK119ub1 is a primary mechanism through which the Polycomb systems mediates transcriptional repression in ESCs.

Interestingly, the contribution of PRC1 and its catalysis to the regulation of mammalian gene expression has recently been examined in the context of neural cell differentiation ^78^. In this case, catalysis was proposed to be necessary for repression during early neurogenesis, although this dependency appeared to be less pronounced in more differentiated cells that were slowly proliferating. This raises the interesting possibility that H2AK119ub1 may be the primary determinant of Polycomb-mediated repression in rapidly proliferating cells where the integrity of Polycomb chromatin domains is constantly challenged by DNA replication. In contrast, other Polycomb chromatin domain features may function to increase the fidelity of gene repression when cells are no longer rapidly dividing and must ensure long-term and stable maintenance of repression. In agreement with this possibility the kinetics of PRC1-dependent H2AK119ub1 addition (unpublished observation) and removal appear to be very rapid, whereas PRC2-dependent H3K27me3 addition and removal is far slower ^79–83^. In the context of these findings, we speculate that PRC1 and H2AK119ub1 could function to rapidly respond to alterations in transcription to initiate repression and that feedback mechanisms that integrate PRC2 activity may function over longer time scales to increase the fidelity and stability of repression.

Although chromatin modifying systems have been widely implicated in controlling gene expression, how they actually regulate the process of transcription has remained poorly understood. Through defining the repressive features of Polycomb chromatin domains and then combining genome-wide measurements of gene expression and Pol II binding with single-molecule single-cell transcription analysis we now gain the first detailed view of the mechanisms that enable Polycomb-mediated gene repression in ESCs. We show that the Polycomb system does not function as an absolute block to transcription. Instead, the majority of cells in the population display at least some expression of Polycomb target genes and low-level binding of Pol II at their promoters. Following PRC1 removal, most cells show moderate increases in Polycomb target gene expression, indicating that the Polycomb system functions to protect against ubiquitous low-level transcriptional signals, and not to actively repress genes that would otherwise become fully transcribed in its absence. Therefore, the Polycomb system appears to play fundamental roles in constantly constraining transcription from lowly expressed gene promoters, explaining the central roles that this system plays in maintaining the fidelity of gene expression patterns and supporting normal cell identity.

These new findings then raise the important question of how the Polycomb system mechanistically counteracts the process of transcription to elicit its repressive effects. By measuring transcription in single cells, we find that Polycomb-dependent gene repression works through regulating transcription burst frequency. While this effect could simply manifest from PRC1/H2AK119ub1 causing local chromatin compaction and formation of higher-order chromatin structures refractory to chromatin remodeling and active transcription as has been proposed previously ^84–88^, we and others have shown that perturbation of the Polycomb system in ESCs does not profoundly influence target gene promoter accessibility ^89–91^. Our interpretation of these observations is that PRC1 and H2AK119ub1 must therefore have more direct roles in counteracting Pol II function. Indeed it has been suggested that the Polycomb system functions to limit the release of a poised or paused polymerase, based on the presence of Ser5 phosphorylated Pol II at Polycomb target gene promoters ^50, 57^. However, evidence from measuring transcriptionally engaged Pol II by genome-wide run-on assays suggests that Polycomb target gene promoters lack paused Pol II ^58, 59^ and we find that derepression of Polycomb target genes requires new transcription initiation, not simply the release of a preexisting engaged polymerase. Furthermore, alterations to transcriptional pause release have been proposed to primarily affect transcription burst sizes ^92, 93^, not frequency as discovered here. Importantly, transcriptional burst frequency can be increased by interaction with enhancers ^94–98^ and this is likely dependent on transcriptional activators that influence Pol II activity at the gene promoter ^99, 100^. Furthermore, histone acetylation has also been implicated in promoting burst frequency when found at gene promoters or enhancers ^98, 101^. Here we find that the Polycomb system elicits the opposite effect on burst frequency and appears to do so through functioning directly at the gene promoter. It is therefore tempting to speculate that the Polycomb system may have evolved to function specifically at gene promoters to counteract other activities that would normally function to increase burst frequency and activate gene transcription. As such, the Polycomb system may play a key role at gene promoters to counteract low level or inappropriate transcription signals that emanate from regulatory elements, like enhancers, that if left unchecked could lead to bursts of transcription that would endanger maintenance of cell identity. Furthermore, we envisage that this Polycomb-dependent repressive activity could also play important roles during the early stages of gene induction to ensure appropriate and persistent activation signals are present before a gene can transition into an activated state, for example, during cell fate transitions ^3, 28^. Therefore, examining the influence of the Polycomb system on transcription dynamics during cellular differentiation will be important in the context of future work.

At a molecular level we envisage that PRC1 and H2AK119ub1 are likely to affect transcription burst frequency by controlling some key aspect of transcription initiation. This is in agreement with previous observations where the Polycomb system was proposed to interfere with the assembly or activity of the pre-initiation complex ^48, 53^. However, defining precisely how H2AK119ub1 influences transcription initiation will require more detailed consideration, and given the complexity of this process, it will be important to use highly defined *in vitro* systems where H2AK119ub1 has been shown to repress transcription ^49, 77^. Nevertheless, our detailed *in vivo* work provides a strong conceptual basis for Polycomb-mediated gene repression which places PRC1 and H2AK119ub1 as central components through which Polycomb chromatin domains control Pol II function and transcriptional burst frequency to regulate gene expression.

## Methods

### List of antibodies

| Antibody | Source | Identifier | Application |
| --- | --- | --- | --- |
| Rabbit monoclonal anti-H2AK119ub1 | Cell Signaling Technology | Cat# 8240 | Western and ChIP |
| Rabbit polyclonal anti-H3K27me3 | In house<sup>71</sup> | n/a | Western and ChIP |
| Rabbit polyclonal anti-H3K4me3 | In house<sup>16</sup> | n/a | ChIP |
| Mouse monoclonal anti-H3 | Cell Signaling Technology | Cat# 3638 | Western |
| Mouse monoclonal anti-H4 | Cell Signaling Technology | Cat# 2935 | Western |
| Rabbit monoclonal anti-RING1B | Cell Signaling Technology | Cat# 5694 | Western and ChIP |
| Mouse monoclonal anti-RING1B | <sup>102</sup> | n/a | IP |
| Rabbit monoclonal anti-SUZ12 | Cell Signaling Technology | Cat# 3737 | Western and ChIP |
| Rabbit monoclonal anti-Rbp1-NTD | Cell Signaling Technology | Cat# 14958 | ChIP |
| Rabbit monoclonal anti-Rbp1-Ser5P | Cell Signaling Technology | Cat# 13523 | ChIP |
| Rabbit monoclonal anti-Rbp1-Ser2P | Cell Signaling Technology | Cat# 13499 | ChIP |
| Rabbit monoclonal anti-RING1A | Cell Signaling Technology | Cat# 13069 | Western |
| Rabbit polyclonal anti-PCGF1 | In house<sup>23</sup> | n/a |  |
| Rabbit polyclonal anti-PCGF2 (Mel-18) | Santa Cruz | Cat# sc-10744 | Western |
| Rabbit monoclonal anti-PCGF3+PCGF5 | Abcam | Cat# ab201510 | Western |
| Rabbit polyclonal anti-PCGF6 | LifeSpan BioSciences | Cat# LS-C482495 | Western |
| Mouse monoclonal anti-PHC1 | Cell Signaling Technology | Cat# 13768 | Western |
| Rabbit polyclonal anti-CBX7 | Millipore | Cat# 07-981 | Western |
| Rabbit polyclonal anti-RYBP (DEDAF) | Millipore | Cat# AB3637 | Western |
| Rabbit polyclonal anti-KDM2B | In house<sup>16</sup> | n/a | Western |
| Mouse monoclonal anti-TBP | Abcam | Cat# ab818 | Western |
| Rabbit polyclonal anti-CFP1 | In house<sup>103</sup> | n/a | Western |

### Cell lines, culturing and treatments

The control TIR1 and PRC1^deg^ cell lines were derived from male E14 mouse embryonic stem cells (ESCs) as described previously ^34^. In brief, the *TIR1* coding sequence from the *Oryza sativa* was inserted into the *Rosa26* locus by CRISPR-Cas9 engineering, generating the TIR1 control line. These cells were further engineered to create the PRC1^deg^ cell line by introducing the full-length auxin inducible degron (AID) tag at the N-terminus of both *Ring1B* alleles, and constitutively deleting *Ring1A* by excising exons 1-3. The PRC2^deg^ cell line was previously generated ^46^ by inserting dTAG at the N-terminus of the endogenous *Suz12* in the *PCGF2-HaloTag* genetic background.

Mouse ESCs were grown on gelatin-coated plates, in Dulbecco’s Modified Eagle Medium (DMEM, Life Technologies) supplemented with 15% fetal bovine serum (FBS, Labtech), 0.5 mM beta-mercaptoethanol (Life Technologies), 2 mM L-glutamine (Life Technologies), 1x penicillin-streptomycin (Life Technologies), 1x non-essential amino acids (Life Technologies) and 10 ng/mL leukemia inhibitory factor. Cells were cultured at 37°C with 5% CO2. For calibration of genomic experiments, we used either *Drosophila* S2 (SG4) cells, cultured at 25°C in Schneider’s Drosophila Medium (Life Technologies), supplemented with 10% heat-inactivated FBS and 1x penicillin-streptomycin, or human HEK293T cells, cultured at 37°C with 5% CO2, in DMEM supplemented with 10% FBS, 2 mM L-glutamine, 1x penicillin-streptomycin and 0.5 mM beta-mercaptoethanol. All cells were regularly tested for the presence of mycoplasma.

To induce AID-RING1B degradation, water-dissolved auxin (indole-3-acetic acid (IAA), Sigma) was mixed with cell medium to the final concentration of 500 µM and added to PRC1^deg^ cells at designated times before harvesting by trypsinisation. To induce dTAG-SUZ12 degradation, PRC2^deg^ cells were treated with 100 nM dTAG-13 compound ^45^ for 2 hours. For the Pol II initiation inhibition experiments, PRC1^deg^ cells were treated with 500 nM triptolide (Merck) or DMSO for 50 minutes before adding auxin for 2 hours, and harvested by scraping in the ice-cold PBS. For retinoic acid (RA) differentiation the cells were processed as outlined in ^104^. Briefly, the cells were plated, allowed to attach and switched to a medium with 1 µM RA and without leukemia inhibitory factor. Cells were grown for 72 hours with medium replaced every 24 hours before proceeding with smRNA-FISH.

### Protein extraction and immunoblotting

To extract nuclear proteins, cell pellets were resuspended in 10 volumes of buffer A (10 mM HEPES pH 7.9, 1.5 mM MgCl2, 10 mM KCl, 0.5 mM DTT, 0.5 mM PMSF, and 1x cOmplete protease inhibitor (PIC, Roche)) and incubated for 10 min on ice. After centrifugation at 1500 g for 5 min, the cell pellets were resuspended in 3 volumes of buffer A supplemented with 0.1% NP-40. After mixing by inversion, nuclei were pelleted again at 1500 g for 5 min. Pelleted nuclei were resuspended in 1 volume of buffer C (250 mM NaCl, 5 mM HEPES pH 7.9, 26% glycerol, 1.5 mM MgCl2, 0.2 mM EDTA, 0.5 mM DTT and 1x PIC). The volume of the nuclear suspension was measured and concentration of NaCl raised to 400 mM by dropwise addition. Nuclei were rotated at 4°C for 1 h to extract nuclear proteins, which were recovered as the supernatant after centrifugation at 18,000 g for 20 min. Protein concentration was determined by Bradford assay (BioRad).

Histone extracts were prepared by washing pelleted cells in RSB (10 mM Tris HCl pH 7.4, 10 mM NaCl, 3 mM MgCl2 and 20 mM N-ethylmaleimide (NEM)), followed by centrifugation at 240 g for 5 min and resuspension in RSB buffer supplemented with 0.5% NP-40. Following incubation on ice for 10 min, cells were centrifuged at 500 g for 5 minutes. After resuspending the nuclear pellet in 5 mM MgCl2, an equal volume of 0.8 M HCl was added, and histones were extracted for 20 min on ice and incubated on ice for 20 min to extract histones. The supernatant was taken after centrifugation for 20 min at 18,000 g, and histones precipitated by addition of TCA to 25% v/v and incubation on ice for 30 min. Histones were pelleted by centrifugation at 18,000 g for 15 min, and washed twice with cold acetone. The histone pellet was resuspended by vortexing in 1x SDS loading buffer (2% SDS, 100 mM Tris pH 6.8, 100 mM DTT, 10% glycerol and 0.1% bromophenol blue) and boiling at 95°C for 5 min. Histones were stained by Coomassie Brilliant Blue following SDS-PAGE to compare concentrations between samples.

For western blot analysis, nuclear or histone extracts were resolved by SDS-PAGE and transferred onto a nitrocellulose membrane using the Trans-Blot Turbo Transfer System (BioRad). Membranes were blocked in 5% milk in 1x PBS with 0.1% Tween 20 (PBST) for at least 30 min at RT. Membranes were incubated in the same buffer with primary antibodies overnight at 4°C, washed 3x 5 min with PBST, and incubated with the IRDye secondary antibodies (LI-COR) in PBST for 60 min at RT. Following 2x 5 min PBST and 1x 5 min PBS wash, membranes were imaged using the Odyssey Fc system (LI-COR). Changes in bulk protein levels were quantified relative to the loading controls (TBP or CFP1 for nuclear extracts, H3 or H4 for histone extracts).

### Co-immunoprecipitation

For co-immunoprecipitation experiments, 500 µg of nuclear extract from the control (TIR1) or PRC1^deg^ cells was diluted to 550 µL with BC150 buffer (150 mM KCl, 10% glycerol, 50 mM HEPES (pH 7.9), 0.5 mM EDTA, 0.5 mM DTT, 1x PIC). A 50 μL aliquot was retained as input, and the rest was incubated overnight at 4°C with 5 μg of mouse monoclonal anti-RING1B antibody ^102^. Protein A agarose beads (Repligen) were used to capture the immunoprecipitated material for 1 h at 4°C. Beads were pelleted at 1000 g, washed three times with BC150, resuspended in 120 μL of 1x SDS loading buffer and boiled at 95°C for 5 min. The supernatant was taken as the immunoprecipitate and analysed by western blotting along with the input, as described above.

### Gene expression analysis by RT-qPCR

RNA was isolated from cell nuclei using either TRIzol reagent (Invitrogen) or RNeasy Mini Kit (Qiagen). Genomic DNA was digested with the TURBO DNA-free Kit (Invitrogen). cDNA was synthesized from 400 ng of RNA using ImProm-II Reverse Transcription system kit and random primers (Promega). Quantitative PCR was performed in technical duplicates using SensiMix SYBR mix (Bioline). The *U6 snRNA* gene was used as an internal control.

### Calibrated nuclear RNA-sequencing (cnRNA-seq)

For cnRNA-seq, 2x10^7^ mouse ESCs (untreated or treated with auxin for indicated times) were mixed with 8x10^6^ *Drosophila* SG4 cells in PBS. Nuclei were released in 1 mL HS Lysis buffer (50 mM KCl, 10 mM MgSO4.7H20, 5 mM HEPES, 0.05% NP40, 1 mM PMSF, 3 mM DTT, 1x PIC) for 1 min at room temperature, and recovered by centrifugation at 1000 g for 5 min at 4°C, followed by three washes with ice-cold RSB buffer (10 mM NaCl, 10 mM Tris pH 7.4, 3 mM MgCl2). Pelleted nuclei were resuspended in 1 mL of TRIzol reagent and RNA was extracted according to the manufacturer’s protocol, followed by treatment with the TURBO DNA-free Kit. Quality of RNA was assessed using 2100 Bioanalyzer RNA 6000 Pico kit (Agilent). Next, rRNA was depleted using the NEBNext rRNA Depletion kit (NEB). RNA-seq libraries were prepared using the NEBNext Ultra Directional RNA Library Prep kit (NEB), indexed using NEBNext Multiplex Oligos (NEB), polled and sequenced as 80 bp paired-end reads on the Illumina NextSeq 500 in biological triplicates.

### Preparation of chromatin for calibrated ChIP-seq (cChIP-seq)

All cChIP-seq experiments were performed in biological triplicates and calibrated using the whole-cell spike-in ^105–107^.

For RING1B and SUZ12, double-crosslinked cChIP-seq was performed as described previously ^30^. Briefly, 5x10^7^ mouse ESCs were first crosslinked with 2 mM disuccinimidyl glutarate (DSG, Thermo Scientific) while rotating for 45 min at 25°C, and then with 1% formaldehyde (methanol-free, Thermo Scientific) for further 15 min. Reactions were quenched by addition of 125 mM glycine. Mouse ESCs were then mixed with 2x10^6^ double-crosslinked HEK293T cells and incubated in lysis buffer (50 mM HEPES pH 7.9, 140 mM NaCl, 1 mM EDTA, 10% glycerol, 0.5% NP-40, 0.25% Triton X-100, 1x PIC) for 10 min at 4°C. The released nuclei were washed (10 mM Tris-HCl pH 8, 200 mM NaCl, 1 mM EDTA, 0.5 mM EGTA, 1x PIC) for 5 min at 4°C. Chromatin was then resuspended in 1 mL of sonication buffer (10 mM Tris-HCl pH 8, 100 mM NaCl, 1 mM EDTA, 0.5 mM EGTA, 0.1% sodium deoxycholate, 0.5% N-lauroylsarcosine, 1x PIC) and sonicated for 30 min using the BioRuptor Pico (Diagenode). Following sonication, Triton X-100 was added to a final concentration of 1%. After centrifugation at 20,000 g for 10 min at 4°C, supernatant was stored at -80°C until use.

For total and phosphorylated Pol II, single-crosslinked cChIP-seq was performed as described previously ^108^. Briefly, 5x10^7^ mouse ESCs were crosslinked in 1% formaldehyde for 10 min at 25°C and then quenched with 125 mM glycine. Mouse ECSs were mixed with 2x10^6^ single-crosslinked HEK293T cells and incubated in FA-lysis buffer (50 mM HEPES pH 7.9, 150 mM NaCl, 2 mM EDTA, 0.5 mM EGTA, 0.5% NP-40, 0.1% sodium deoxycholate, 0.1% SDS, 10 mM NaF, 1 mM AEBSF, 1x PIC) for 10 min.

Chromatin was sonicated for 30 min using the BioRuptor Pico, centrifuged at 20,000 g for 10 min at 4°C, and supernatant was stored at -80°C until use.

For H2AK119ub1, H3K27me3 and H3K4me3, native cChIP-seq was performed as described previously^30^. Briefly, 5x10^7^ mouse ESCs were mixed with 2x10^7^ *Drosophila* SG4 cells in PBS. The cells were pelleted and nuclei were released by resuspending in ice-cold lysis buffer (10 mM Tris-HCl pH 8, 10 mM NaCl, 3 mM MgCl2, 0.1% NP-40, 5 mM NEM). Nuclei were then washed, and resuspended in 1 mL of MNase digestion buffer (10 mM Tris-HCl pH 8, 10 mM NaCl, 3 mM MgCl2, 0.25 M sucrose, 3 mM CaCl2, 10 mM NEM, 1x PIC). Samples were digested with 150 units of MNase (Fermentas) for 5 min at 37°C, stopped by 4 mM EDTA. The supernatant (S1) was collected following centrifugation at 1500 g for 5 min at 4°C. The remaining pellet was incubated with 300 μl of nucleosome release buffer (10 mM Tris-HCl pH 7.5, 10 mM NaCl, 0.2 mM EDTA, 10 mM NEM, 1x PIC) at 4°C for 1 h, passed five times through a 27G needle, and spun at 1500 g for 5 min at 4°C. The second supernatant (S2) was collected and combined with corresponding S1 supernatant, and stored at -80°C until use. Digestion to predominantly mono-nucleosomal fragments was confirmed by agarose gel electrophoresies of purified DNA.

### Chromatin immunoprecipitation and massively parallel sequencing

For double-crosslinked ChIP, sonicated chromatin was diluted 10-fold in ChIP dilution buffer (1% Triton X-100, 1 mM EDTA, 20 mM Tris-HCl pH 8, 150 mM NaCl, 1x PIC) and pre-cleared for 1 h with either protein A agarose beads (Repligen, for RING1B ChIP) or protein A magnetic Dynabeads (Invitrogen, for SUZ12 ChIP) blocked with 1 mg/mL bovine serum albumin (BSA) and 1 mg/mL yeast tRNA. For each ChIP reaction, 1 mL of diluted and pre-cleared chromatin was incubated overnight with the appropriate antibody, either anti-RING1B (CST, D22F2, 3 μl), or anti-SUZ12 (CST, D39F6, 3 μl). Antibody-bound chromatin was captured using blocked protein A beads (agarose for RING1B, magnetic Dynabeads for SUZ12) for at least 2 h at 4°C and collected by centrifugation/on a magnetic rack. ChIP washes were performed as described previously ^16^. ChIP DNA was eluted in elution buffer (1% SDS, 0.1 M NaHCO3) and cross-linking was reversed overnight at 65°C with 200 mM NaCl and 2 μL RNase A (Sigma). A matched input sample (1/10 of original ChIP reaction) was treated identically. The following day, ChIP samples and inputs were incubated with Proteinase K (Sigma) for 1.5 h at 56°C and purified using ChIP DNA Clean and Concentrator Kit (Zymo Research).

For Pol II ChIP, 300 ug of chromatin was diluted to 1 ml in FA-lysis buffer and pre-cleared for 1 h with protein A agarose beads blocked with 1 mg/mL BSA and 1 mg/mL yeast tRNA. For each ChIP reaction, diluted and pre-cleared chromatin was incubated overnight with the appropriate antibody, anti-Rbp1-NTD (CST, D8L4Y, 15 μl) to detect total Pol II, anti-Rbp1-CTD-Ser5P (CST, D9N5I, 12.5 μl), or anti-Rbp1-CTD-Ser2P (CST, E1Z3G, 12.5 μl). Antibody-bound chromatin was isolated using blocked protein A agarose beads for 3 h at 4°C. Washes were performed with FA-Lysis buffer, FA-Lysis buffer containing 500 mM NaCl, DOC buffer (250 mM LiCl, 0.5% NP-40, 0.5% sodium deoxycholate, 2 mM EDTA, 10 mM Tris–HCl pH 8), followed by two washes with TE buffer pH 8. ChIP DNA was eluted, de-crosslinked and purified as described above, along with the matched input samples.

Native chromatin was diluted 10-fold in native ChIP incubation buffer (70 mM NaCl, 10 mM Tris–HCl pH 7.5, 2 mM MgCl2, 2 mM EDTA, 0.1% TritonX-100, 10 mM NEM, 1x PIC). For each ChIP reaction, 1 ml of diluted nucleosomes was incubated overnight at 4°C with the appropriate antibody, anti-H2AK119ub1 (CST, D27C4, 5 μL), anti-H3K27me3 (in-house, 5 μL) or anti-H3K4me3 (in-house, 3 μL). Antibody-bound nucleosomes were captured by incubation for 1 h at 4°C with protein A agarose beads, pre-blocked overnight in native ChIP incubation buffer supplemented with 1 mg/ml BSA and 1 mg/ml yeast tRNA. The beads were then washed four times with native ChIP wash buffer (20 mM Tris– HCl pH 7.5, 2 mM EDTA, 125 mM NaCl, 0.1% TritonX-100) and once with TE buffer pH 8. Immunoprecipitated DNA was eluted using 100 μl of elution buffer (1% SDS, 0.1 M NaHCO3) and purified using ChIP DNA Clean and Concentrator Kit. DNA from a matched input sample (corresponding to 10% of the original ChIP reaction) purified in the same way.

cChIP-seq libraries for both ChIP and input samples were prepared using NEBNext Ultra DNA Library Prep Kit for Illumina (double-crosslinked and native ChIP-seq) or NEBNext Ultra II DNA Library Prep Kit for Illumina (Pol II ChIP-seq), following manufacturer’s guidelines. Samples were indexed using NEBNext Multiplex Oligos. The average size and concentration of all libraries was analysed using the 2100 Bioanalyzer High Sensitivity DNA Kit (Agilent) followed by qPCR using SensiMix SYBR (Bioline, UK) and KAPA Illumina DNA standards (Roche). Libraries were sequenced as 40 bp paired-end reads on the Illumina NextSeq 500.

### Single molecule RNA fluorescence in situ hybridization

smRNA-FISH probe sets comprised 48 individually labelled oligos that were 20-22nt in length and designed using the online tool from the manufacturer (Stellaris). Mouse ESCs were harvested at designated times after treatment with IAA, fixed with 3.7 % formaldehyde for 10 min and then incubated in 70% (v/v) ethanol/PBS solution for at least 1 hour at 4°C. smRNA-FISH was carried out in suspension following the standard guidelines from the manufacturer (Stellaris). Briefly, the cells were pelleted and resuspended in buffer containing 10% formamide and 2x SSC (buffer A), followed by centrifugation at 300 g and further resuspension in hybridisation mixture comprising 20% dextran sulfate, 10% formamide, 2x SSC and probe sets specific to exons (conjugated with Q570) or introns (with Q670), in a total volume of 200 µl per cell pellet. After overnight incubation at 37°C, the cells were washed twice with buffer A. DNA was stained with DAPI dissolved in buffer A and cell membranes with Agglutinin conjugated with Alexa 488 (Thermofisher) in PBS. After a final wash and PBS aspiration the cell pellet’s volume was approximately 150 µl; 6 µl of the pellet was mixed with 10 µl of Vectashield (H-1000, Vectorlabs) and spread on a glass slide. The solution was covered with microscopy coverslip and pressed to remove excess mounting medium and ensure cell monolayer.

Polycomb target genes for study by smRNA-FISH were chosen based on transcript information from cnRNA-seq and fulfilled following criteria: they spanned a wide range of basal expression levels and had varying magnitude of derepression upon RING1B depletion (Figure S7A). Control genes (*Tfrc* and *Hspg2*) had moderate expression levels and were classified as non-PcG (see below). Whole-cell smRNA-FISH results were cross-validated with cnRNA-seq and were mostly in good agreement (Figure S7B).

### Microscopy

Microscopy was carried out using Olympus IX-83 system and Cell Sens software, equipped with 63x1.4 NA oil objective lens and 1200x1200px^2^ sCMOS camera (Photometrics), 91.5 nm pixel size. Fluorescence was excited with LED light (Coolled): 365 nm for DAPI, 488 nm for Alexa 488, 550 nm for Q570, and 635 nm for Q670. 3D-stacks contained 30 images and were acquired every 350 nm. Typically, 40 to 60 fields of view were acquired per sample, yielding between 1500 – 3000 cells in total from 3 biological replicates.

### Quantification and Statistical Analysis Sequencing data processing and normalization

For cnRNA-seq, paired-end reads were first aligned using Bowtie 2 (“–very-fast,” “–no-mixed” and “– no-discordant” options) ^109^ against the concatenated mm10 and dm6 rDNA genomic sequence (GenBank: BK000964.3 and M21017.1) in order to discard the reads mapping to rDNA. The remaining unmapped reads were then aligned against the concatenated mm10 and dm6 genome using STAR ^110^. Reads which failed to map using STAR were further aligned against the concatenated mm10 and dm6 genome using Bowtie 2 (“–sensitive-local,” “–no-mixed” and “–no-discordant” options), to improve the overall mapping of intronic sequences of nascent transcripts present in nuclear RNA fraction. Uniquely mapped reads from the two alignment steps were combined and PCR duplicates were removed using Sambamba ^111^ before the further analysis.

For cChIP-seq, paired-end reads were aligned to the concatenated mouse (mm10) and spike-in (dm6 for native, hg19 for cross-linked cChIP-seq) genome sequences using Bowtie 2 (“–no-mixed” and “– no-discordant” options). Only uniquely mapped reads were kept for downstream analysis, after removal of PCR duplicates with Sambamba. All cnRNA-seq and cChIP-seq experiments performed in this study are listed in Table S1 along with the number of uniquely aligned reads for both mouse and spike-in genomes.

For visualization of cChIP-seq sequencing data and annotation of genomic regions with read counts, uniquely mapped mouse reads were normalized using dm6 or hg19 spike-in as described previously ^30^. Briefly, mouse reads were randomly subsampled based on the total number of spike-in (dm6 or hg19) reads in each sample. Additionally, to account for possible minor variations in spike-in cell mixing between cChIP-seq replicates, we corrected the subsampling factors by using the ratio of spike-in to mouse total read counts in the corresponding input samples. Prior to merging normalized replicates for visualization and analysis, read coverage across regions of interest (RING1B peaks for RING1B, SUZ12, H2AK119ub1 and H3K27me3, or TSS ± 2.5 kb for total Pol II, Ser5P-Pol II and H3K4me3, or gene bodies for Ser2P) was analyzed using multiBamSummary and plotCorrelation functions from deepTools (v3.1.1) ^112^, confirming a high correlation between replicates (Pearson’s correlation coefficient > 0.9) (Table S2). Genome coverage tracks for cChIP-seq were generated using the pileup function from MACS2 ^113^ and visualized using the UCSC genome browser ^114^. Differential genome coverage tracks (log2 ratio of the two conditions) were made by using bigwigCompare from deeptools.

### Peak calling

To identify genomic regions bound by PRC1 we performed peak calling using MACS2 (“BAMPE” and “–broad” options) on the RING1B cChIP-seq data with corresponding input samples for background normalization. A set of peaks identified in all biological replicates of untreated PRC1^deg^ cells was used for further analysis, after filtering out several peaks identified following 4 hours of auxin treatment (i.e. following loss of RING1B) and manually removing sequencing artefacts. In total, 7240 stringent RING1B peaks were identified.

### Differential gene expression analysis and gene annotation

For differential gene expression analysis, we obtained read counts from the original bam files prior to spike-in normalization for a non-redundant mouse gene set using a SAMtools-based custom Perl script. The non-redundant mm10 gene set (n = 20633) was derived from mm10 refGene genes by filtering out very short genes with poor sequence mappability and highly similar transcripts. To identify significant gene expression changes following auxin treatment, we used a custom R script that incorporates spike-in calibration into DESeq2 analysis ^115^ and uses “apeglm” method for LFC shrinkage^116^. Spike-in calibration was incorporated by using read counts from a set of unique dm6 refGene genes to calculate DESeq2 size factors for normalization of raw mm10 read counts, as previously described in ^30, 117^. For a change to be considered significant, we applied a threshold of p-adj < 0.05 and fold change > 1.5. Based on the earliest time point when these thresholds were reached, derepressed genes were classified into three groups (“2 hours”, “4 hours” and “8 hours”). For visualization purposes, DESeq2-normalized read counts were averaged across the replicates and used to calculate RPKM. Log2 fold change values were visualized with MA plots, heatmaps and boxplots made using R and ggplot2. Boxes for the boxplots show interquartile range (IQR), center line represents median, and whiskers extend by 1.5x IQR. Correlation between log2 fold changes in expression between PRC1^deg^ and PRC1^cko^ cells was expressed as Pearson correlation coefficient and visualized with ggplot2 using scatterplot colored by density with stat_density2d. Linear regression line was plotted using geom_smooth along with the R^2^ coefficient of determination.

Mouse genes from the custom non-redundant set (n = 20633) that was used for differential gene expression analysis were classified into three groups based on the overlap of their promoters (TSS ± 2500 bp) with RING1B-bound sites and non-methylated CpG islands (NMI). NMIs are regions enriched in non-methylated CpG dinucleotides, identified using MACS2 peak calling on BioCAP-seq data (n = 27047) ^118^. Genes with promoters that do not overlap NMIs were classified as non-NMI (n = 5534). The rest of the genes were defined as Polycomb-occupied genes (PcG, n = 6937) if their promoters overlapped RINGB-bound sites identified in this study, and those that did not overlap as non-PcG genes (n = 8162). For genes with several alternative transcripts and promoters, we assigned all transcripts into the PcG group if at least one promoter overlapped a RINGB-bound site.

### Read count quantitation and enrichment analysis for cChIP-seq

Metaplot and heatmap analysis of cChIP-seq read density at regions of interest was performed with computeMatrix and plotProfile/plotHeatmap from deepTools ^112^. Metaplot profiles represented mean scores over sets of genomic regions, unless stated otherwise. Read coverages for genomic regions of interest from merged spike-in normalized replicates were computed with multiBamSummary from deeptools (“–outRawCounts”), and used for comparative boxplot analysis. To characterize low-level genomic blanket of H2AK119ub1, we counted reads in a set of 100 kb windows spanning the genome (n = 27282) defined by using the makewindows function from BEDtools (v2.17.0). RPKM values were calculated by dividing normalized read counts by a genomic interval size in kb and visualized with boxplots using custom R scripts and ggplot2. For boxplots comparing H2AK119ub1 cChIP-seq signal at RING1B peaks and 100 kb genomic windows, RPKM values are shown relative to the median signal at RING1B peaks in untreated cells.

Differential enrichment analysis of cChIP-seq data was performed similarly to the gene expression analysis described above, with a few differences. Namely, read counts from individual biological replicates prior to spike-in normalization were obtained with multiBamSummary from deeptools (“– outRawCounts”). Mouse reads were counted in the promoter regions (TSS ± 2.5 kb) of the custom non-redundant mm10 gene set, and over whole gene bodies (TSS to TES) for the analysis of total Pol II. Spike-in genomic reads were counted in the appropriate control regions: human (hg19) CpG islands obtained from UCSC Table Browser for RING1B and SUZ12, human (hg19) promoter regions (TSS ± 2.5 kb) of a custom unique gene set for Pol II, fly (dm6) promoter regions (TSS ± 2 kb) of a custom unique gene set for H3K4me3. Prior to quantification, spike-in reads were pre-normalized using the spike-in/mouse read ratio derived from the corresponding input sample in order to account for minor variations in spike-in cells mixing. For a change to be called significant, we applied a threshold of p-adj < 0.05 and fold change > 1.5.

### Image Analysis

3D images were pre-processed using a custom ImageJ script which we call ThunderFISH. In brief, 3D smRNA-FISH images containing sparse transcripts appearing as diffraction limited spots were converted into 2D images through maximal projection. Due to low numbers of transcripts per cell this operation did not significantly affect the number of transcripts detected when compared with other spot counting methods relying on 3D detection (Figure S6D, E). Slight transcript undercounting was identified only in cells with more than approximately 70 transcripts/cell (Figure S6D). In contrast, most of the samples we imaged contained on average only a few transcripts per cell. In parallel, a threshold was applied on 2D average projections of DAPI and Agglutinin channels, and after converting into binary image and water-shedding, single cell masks were obtained (Figure S6A). The masks with low circularity, touching image boarders, and outside of a specified size range were discarded. Next, the remaining cell masks were used to extract single cell 2D smRNA-FISH images from the full field of view. Lastly, single cell images were converted to a data stack with each frame representing a single 2D smRNA-FISH image of an individual cell. This stack was used as input for diffraction-limited spot counting by the dedicated ImageJ plugin ThunderSTORM ^119^. We used default settings, except for the threshold factor, which we adjusted to be 6-10x the background standard deviation. This range of values provided comparable transcripts per cell distributions (Figure S6B), hence we used it for our transcript detection. ThunderSTORM outputs the number of spots detected per frame (per cell). We used these numbers to produce density graphs of the transcript distributions using R and the ggplot2 package. The pre-processing by ThunderFISH allowed increased imaging and analysis throughput of up to 10,000 cells a day. This proved essential for accurately and efficiently counting Polycomb target gene transcripts, which are lowly expressed, and for extracting robust measures of expression change when the Polycomb system was disrupted.

Nascent RNA-FISH data were analysed using a custom-made ImageJ script utilizing 3D Objects counter^120^ that outputs number of nascent spots per cell with respective integrated intensity.

### Two-state model

All transcript distributions were positively skewed and over-dispersed with a variance at least 2x larger than a mean, and poorly fitted a Poisson distribution. In order to extract inferred mean transcription burst size (number of transcripts produced in a single transcription event), the transcript distribution in a cell population was approximated using a negative binomial distribution fit using the vcd package in R. This distribution is expected when the probability of burst initiation is constant in time and number of transcripts produced per burst is expected to follow exponential distribution. The mean burst size was calculated as *b = (1-p)/p*, where *p* denotes probability of promoter transition to the OFF state extracted from the individual distribution Maximum Likelihood estimation, similarly to as has been described previously ^101, 121^. The goodness of fit of the negative binomial distribution estimated p-values (χ^2^ test) in the majority of the cases exceeded 0.1 (Figure S8B) and were many orders of magnitude greater than if a Poisson distribution was assumed. Therefore, a negative binomial distribution was a better fit to the smRNA-FISH transcript-count distributions.

The frequency of transcription bursts, kON, for individual transcript distributions was computed using following equation ^122^: kON = µƐ/b, where µ represents mean number of transcripts per cell, Ɛ corresponds to transcript half-life, and b corresponds to transcription burst size obtained from negative binomial fit. Since fold changes in kON were computed, transcript life-time Ɛ cancelled out as it was assumed constant and independent of auxin treatment.

### Decomposition of sources of transcript heterogeneity

The two-state model assumes that transcript variability merely stems from an intrinsic stochastic nature of the studied promoter. In order to prove validity of this assumption we sought assessment of extrinsic sources of heterogeneity through decomposing overall transcript distribution heterogeneity. Previously it has been demonstrated that apart from intrinsic behaviour of the promoter, cell cycle and cell volume contribute most to the transcript heterogeneity in a cell population ^123^. We used the previously published formula for cell volume-corrected noise measure *N* ^124^ to estimate the contribution of cell area (proxy for cell volume) and DAPI fluorescence intensity (proxy for cell cycle) to overall measured noise expressed as squared coefficient of variation of transcript distribution CV^2^:

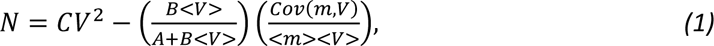

where <V> represents either average area or DAPI intensity, <M> is average number of transcripts, *A* and *B* correspond to the intercept and slope of a linear fit, respectively, of which the best line-fit for the relationship of transcripts and tested extrinsic parameter. Cov(m, V) stands for covariance between number of transcripts *m* and tested extrinsic parameter V. We note that for the most of the genes the linear slope *B* oscillates around 0, effectively cancelling the expression on the right side of the equation 1. Moreover, we note that cell area and DAPI intensity correlated well (Pearson’s r = 0.74, Figure S9B), likely both representing to some extent the same extrinsic cell parameter (likely the cell volume). Nucleus volume has been reported to increase in G2 ^124^, and since the nucleus constitutes the majority of the mouse ESC cell volume, correlation between DAPI intensity and total cell volume in ESCs can be expected, opposite to what was previously reported for another cell type ^124^. Hence, our overall extrinsic noise estimation in spite of yielding miniscule values, is likely still a subject to overestimation. Nevertheless, intrinsic noise for all Polycomb target genes dominated total noise by >90%, and for vast majority of genes almost entirely (Figure S9C). Hence, raw transcript-per-cell distributions were used to infer transcription kinetics based on the two-state model.

### Data and Software Availability

The high-throughput sequencing datasets generated for this study are available in GEO database under the accession number GSEXXXXXX. Published data used in this study include BioCAP-seq (GEO: GSE43512) from ^118^ and cnRNA-seq in *Ring1a^-/-^; Ring1b^fl/fl^* (PRC1^cko^)(GEO: GSE119619) from ^30^. All R and Perl scripts used for genomic data analysis in this study are available upon request. A custom-made ImageJ script for pre-processing 3D images (ThunderFISH) is publicly available with detailed manual of sample preparation and script usage at https://github.com/aleks-szczure/ThunderFISH.

## Acknowledgements

Work in the Klose lab is supported by the Wellcome Trust (209400/Z/17/Z), the European Research Council (681440) and the Lister Institute of Preventive Medicine. We are grateful to Amanda Williams at the Department of Zoology, University of Oxford, for sequencing support on the NextSeq 500. We express our gratitude to Darragh Ennis and Ilan Davis for their advice in RNA-FISH probe design. We thank Nadezda Fursova for her valuable advice on computational analysis. We are grateful to Nadezda Fursova, Neil Blackledge and Emilia Dimitrova for critical reading of the manuscript.

## Author contributions

Conceptualization, P.D., A.S. and R.J.K.; Methodology, Investigation and Formal Analysis, P.D. and A.S.; Writing, P.D., A.S. and R.J.K; Funding Acquisition and Supervision, R.J.K.

---

**Supplementary Figure 1.**
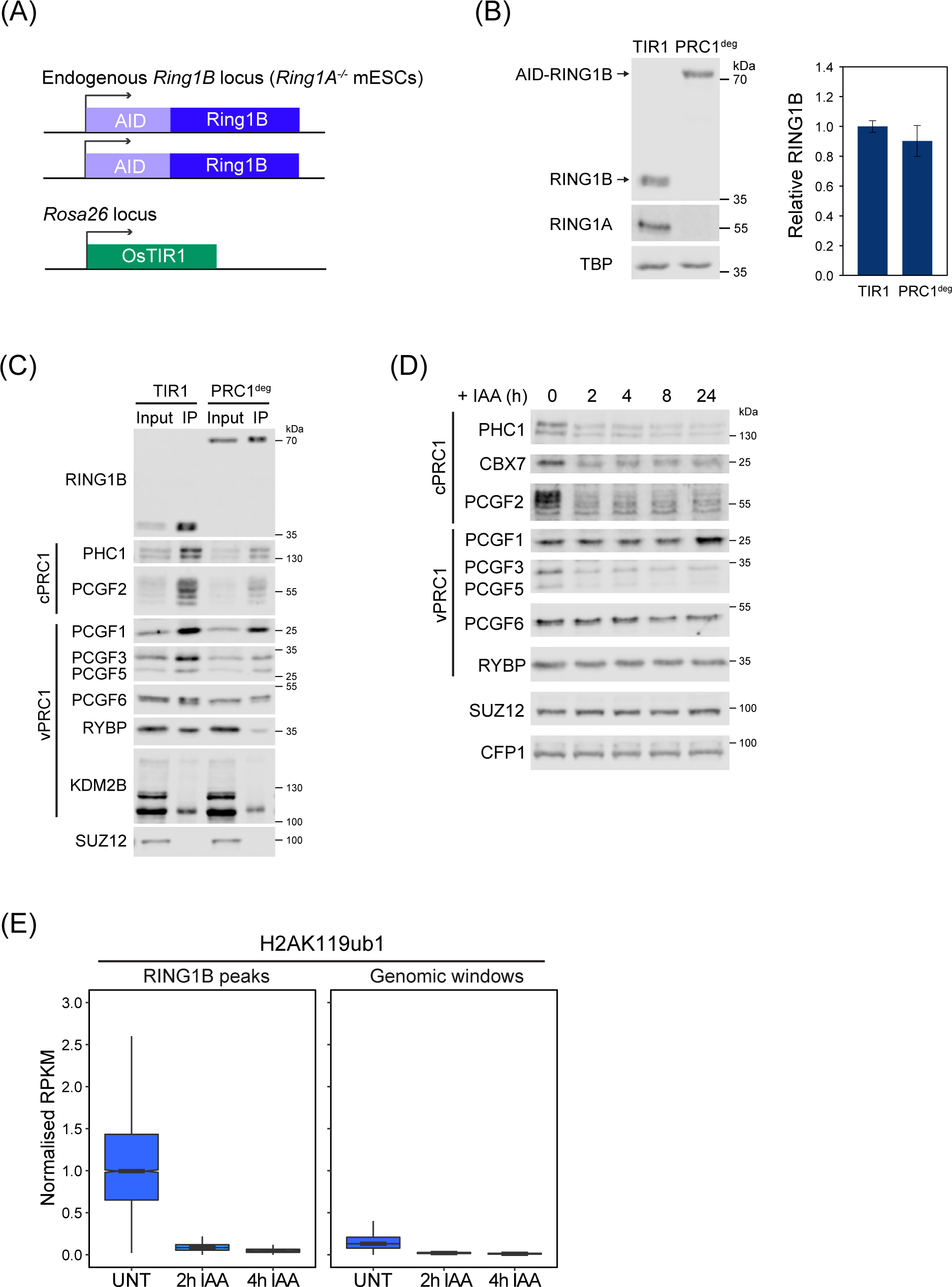
(A) Schematic of the PRC1^deg^ system. Both endogenous *Ring1B* alleles are N-terminally fused to the full-length AID tag in a *Ring1A^-/-^* background, while *OsTIR1* is inserted in *Rosa26* locus. (B) Western blot analysis of RING1A and RING1B in the control cell line (TIR1) and PRC1^deg^ cells (left panel) with a quantification of RING1B levels (right panel). Data represents mean (n=2) ± SD. (C) Immunoprecipitation of RING1B from TIR1 and PRC1^deg^ nuclear extracts followed by western blot analysis of PRC1 components, and SUZ12 (PRC2 component) as a negative control. Shown are representative results from the two independent experiments. vPRC1 – variant PRC1 components, cPRC1 – canonical PRC1 components. (D) Western blot analysis of PRC1 components and SUZ12 in the nuclear extracts of the PRC1^deg^ cells following IAA treatment. CFP1 serves as a loading control. Shown are representative results from the two independent experiments. (E) Boxplots comparing H2AK119ub1 cChIP-seq signal before and after IAA treatment at RING1B-bound sites and over 100 kb genomic windows. All signal is normalised to the median RPKM value of RING1B-bound sites in untreated cells.

**Supplementary Figure 2.**
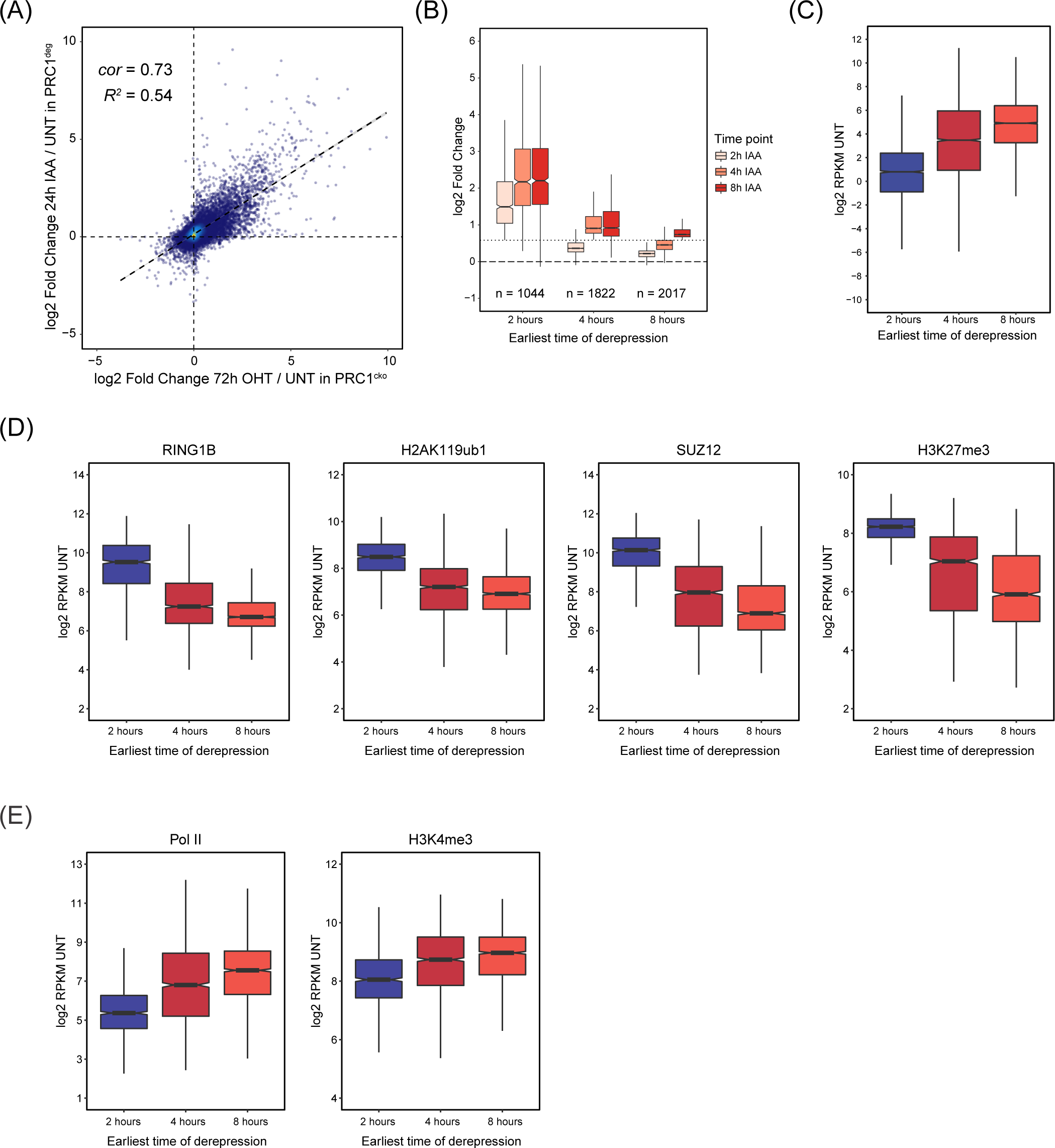
(A) A scatterplot comparing the log2 fold changes in gene expression (cnRNA-seq) in PRC1^deg^ (24h IAA) and PRC1^cko^ (72h OHT) ESCs. *R^2^* is the coefficient of determination for linear regression and *cor* is Pearson correlation coefficient. (B) Boxplots comparing the log2 fold changes in expression of genes split into three groups based on the earliest time point when they became derepressed. Dotted line represents 1.5 fold change. (C) Boxplots comparing the gene expression levels in the untreated PRC1^deg^ cells for the three groups of derepressed genes defined in (B). (D) Boxplots comparing the promoter (TSS ±2.5 kb) cChIP-seq signal for RING1B, H2AK119ub1, SUZ12 and H3K27me3 in the untreated PRC1^deg^ cells for the three groups of derepressed genes defined in (B). (E) Boxplots comparing the promoter (TSS ± 2.5 kb) cChIP-seq signal for total Pol II and H3K4me3 in the untreated PRC1^deg^ cells for the three groups of derepressed genes defined in (B).

**Supplementary Figure 3.**
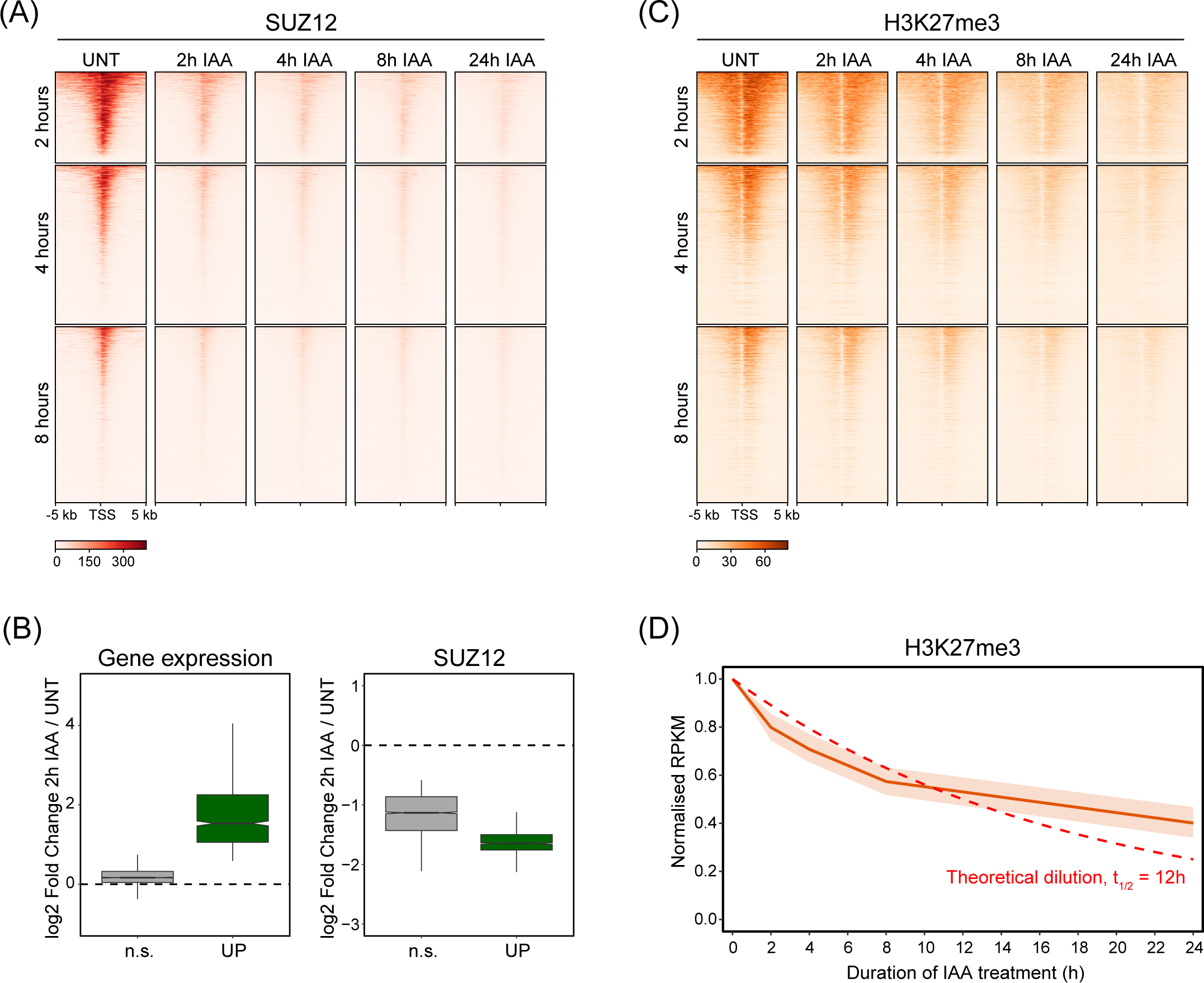
(A) Heatmap analysis of SUZ12 cChIP-seq at TSSs for the three groups of genes defined by the earliest time of derepression in PRC1^deg^ cells treated with IAA for the indicated times. Heatmaps are sorted by RING1B cChIP-seq signal in untreated cells. (B) Box plots illustrating log2 fold changes in expression (cnRNA-seq, left panel) and SUZ12 cChIP-seq signal (right panel) at promoters (TSS ± 2.5 kb) of Polycomb target genes showing a significant reduction in SUZ12 levels following 2 hours of IAA treatment. Genes are divided into Polycomb target genes that become derepressed (UP, n = 955) and those that do not change in expression (n.s., n = 3739) by 2 hours. (C) As in (A) but for H3K27me3 cChIP-seq. (D) The dynamics of reduction in H3K27me3 cChIP-seq signal at RING1B-bound sites which show a significant reduction in H3K27me3 levels by 24 hours of IAA treatment (n = 5926). A theoretical exponential decay function is shown in red, assuming that H3K27me3 levels are halved with every cell cycle if maintenance is completely disrupted. The doubling time of mouse ESCs is approximately 12 hours.

**Supplementary Figure 4.**
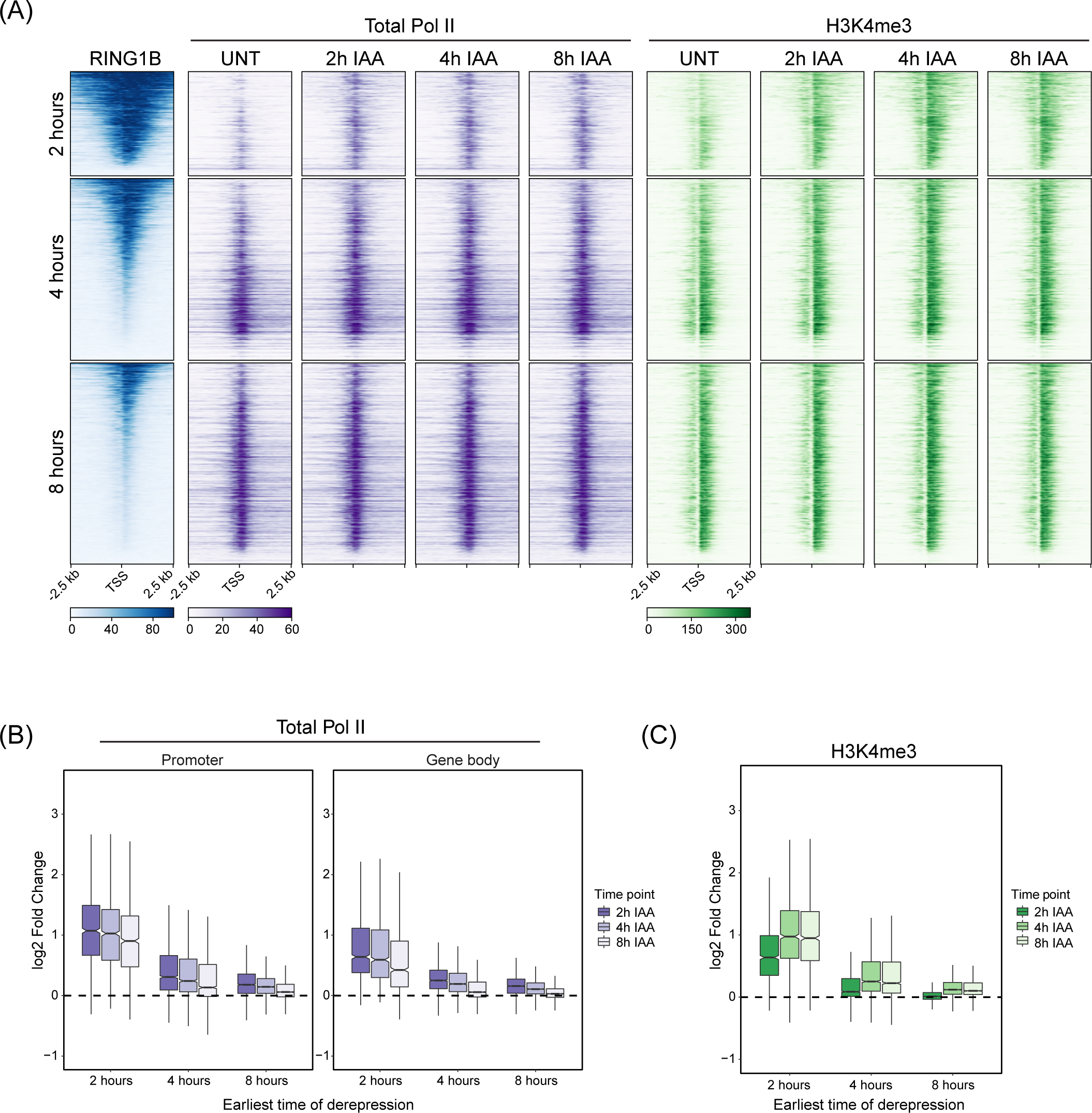
(A) Heatmaps illustrating RING1B binding in untreated cells, and total Pol II and H3K4me3 cChIP-seq signal at TSSs of the three groups of genes defined by the earliest time of derepression in PRC1^deg^ cells treated with IAA for the indicated times. Heatmaps are sorted by RING1B signal in untreated cells. (B) Box plots comparing the log2 fold changes in total Pol II cChIP-seq signal following IAA treatment for genes split into three groups defined by the earliest time of derepression. The analysis was done at promoters (TSS ± 2.5 kb) and over gene bodies (TSS to TES). (C) Box plots comparing the log2 fold changes in H3K4me3 cChIP-seq signal following IAA treatment at promoter regions (TSS ± 2.5 kb) for genes split into three groups defined by the earliest time of derepression.

**Supplementary Figure 5.**
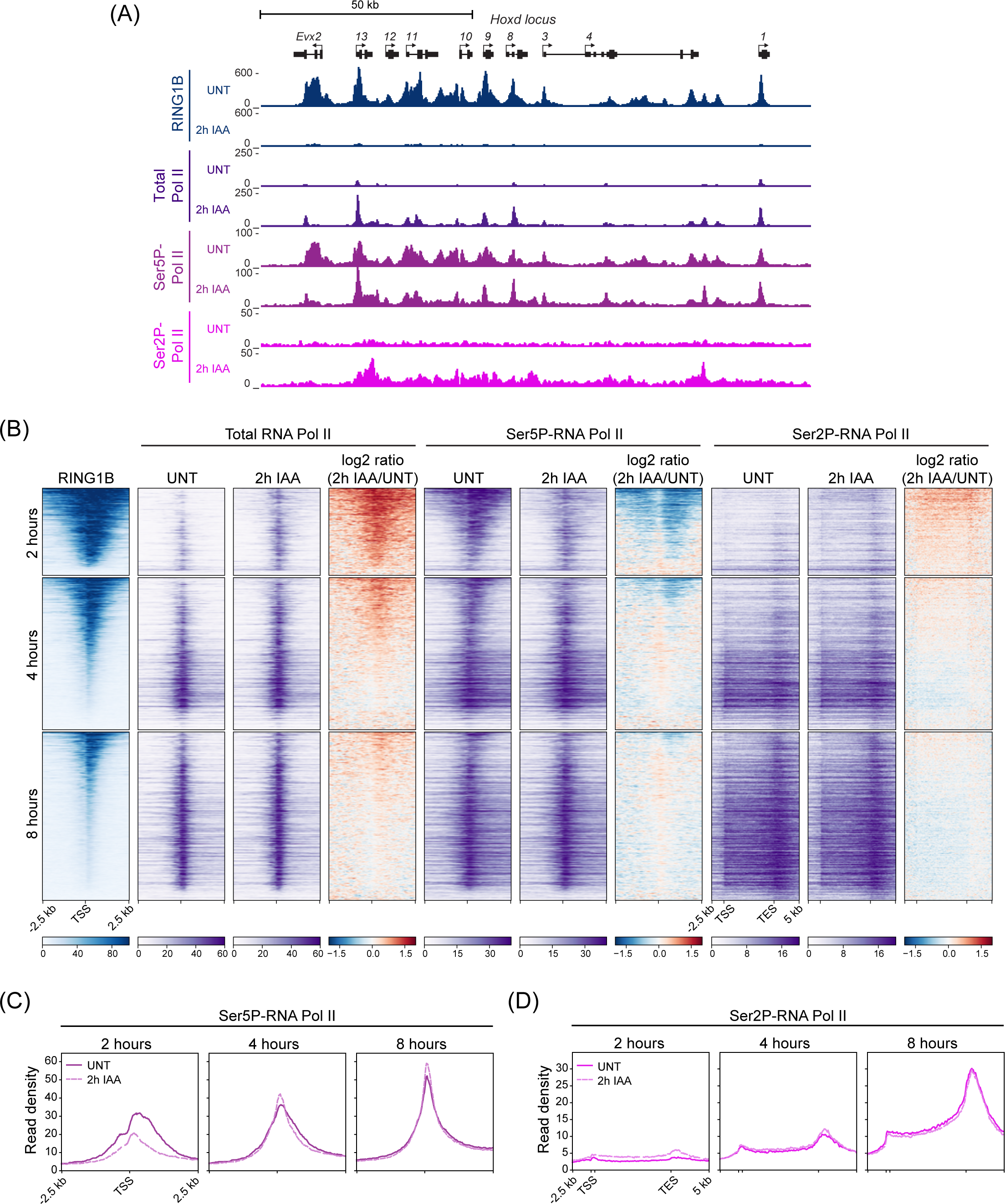
(A) A genomic snapshot of *Hoxd* locus showing cChIP-seq signal for RING1B, total Pol II, Ser5P-Pol II and Ser2P-Pol II in untreated PRC1^deg^ cells and 2 hours after IAA addition. (B) Heatmaps illustrating RING1B, total Pol II, Ser5P-Pol II and Ser2P-Pol II levels and changes in polymerase occupancy following 2h of IAA treatment for three groups of genes defined by the earliest time of derepression. Heatmaps are sorted by RING1B signal in untreated cells. (C) Metaplot analysis of Ser5P-Pol II cChIP-seq at promoters of the three groups of genes defined by the earliest time of derepression in untreated PRC1^deg^ cells and 2 hours after IAA addition. The profiles represent the median signal over the shown genomic region. (D) As in (C) but for Ser2P-Pol II cChIP-seq over gene bodies.

**Supplementary Figure 6.**
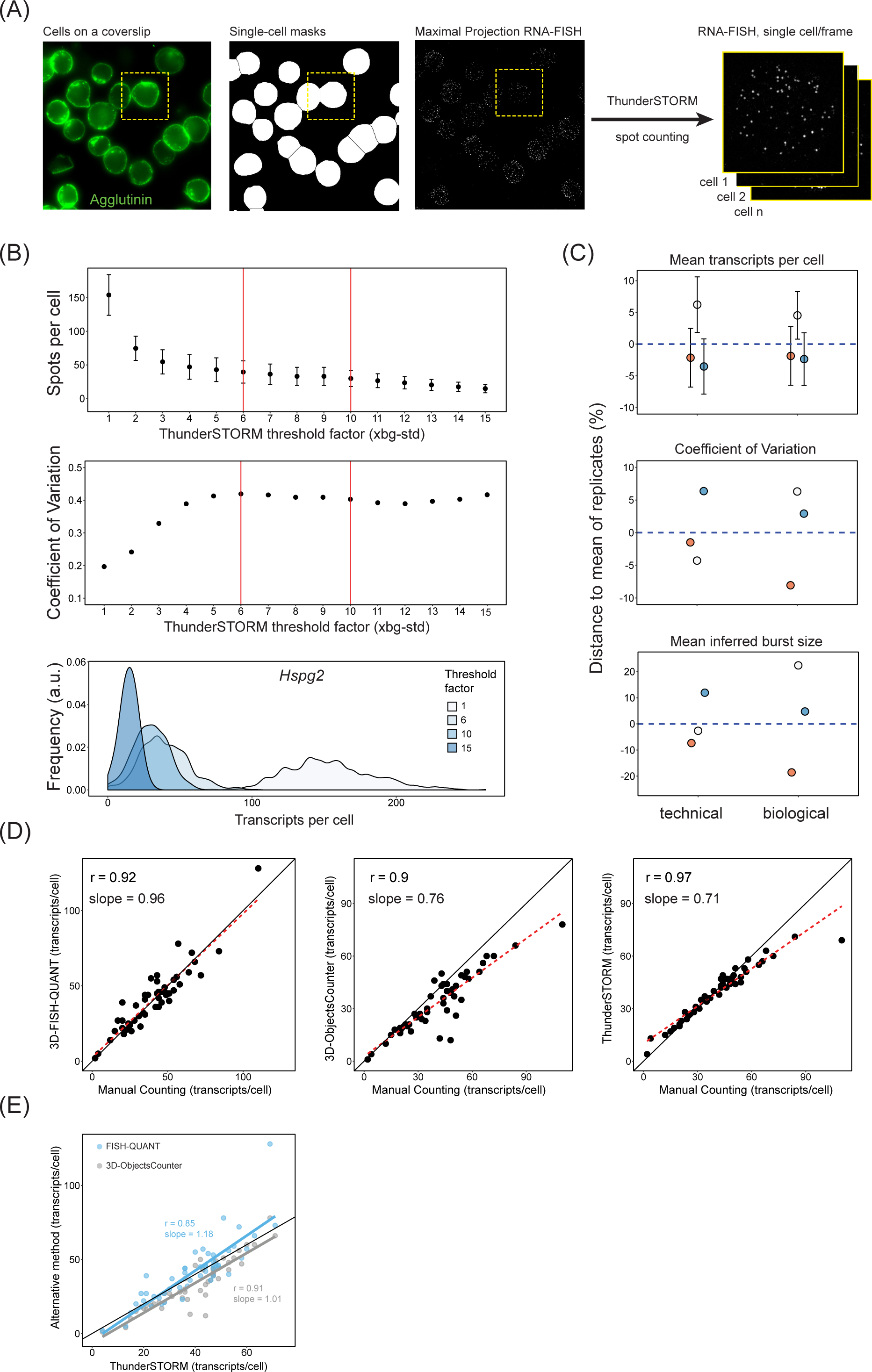
A) A schematic illustrating our automated approach to analyse smRNA-FISH in a high throughput manner. This approach enables effective single-cell segmentation, conversion of the field of view into single-cell smRNA-FISH images, in which individual transcripts can then be counted using ThunderSTORM ^119^. B) An illustration of the effectiveness of ThunderSTORM threshold factor (TF) value identification for spot (transcript) detection in cells. An *Hspg2* dataset is shown as an example. The TF unit is set at a standard deviation of the image background. Spots per cell (top) and heterogeneity of detected transcript levels expressed as coefficient of variation (middle) are shown for a range of TF values. Vertical red lines indicate the range of TF values yielding similar spot-counting outcomes and which were ultimately employed in this study. Density plots (bottom) demonstrate transcript per cell distributions depending on the TF used. TFs from 6 to 10 yield comparable spot-counting values. Very large or very low TF values lead either to overcounting or undercounting of transcript signals. C) An illustration of the reproducibility between technical and biological replicates for our smRNA-FISH transcript counting approach. Mean transcripts per cell (top), coefficient of variation (middle), and mean transcription burst size inferred from the 2-state model (bottom) are shown. Error bars in the top panel represent 10% of standard deviation of transcripts per cell distributions. D) To ensure the robustness of our transcript counting approach, we compared it to other spot (transcript) counting methods and manual counting of transcripts in 50 cells. The methods compared are: 3D-FISH QUANT ^125^ (left), 3D Objects Counter ^120^ (middle), and technique used in this study (right). The right panel indicates that our technique can be prone to a slight undercounting when number of transcripts per cell exceeds 60-70, but otherwise performs comparably to other approaches. Pearson correlation coefficient (r) and slope derived from linear regression are presented. E) Transcript counting using 3D-FISH Quant and 3D Objects Counter correlate well with the transcripts counting using our approach.

**Supplementary Figure 7.**
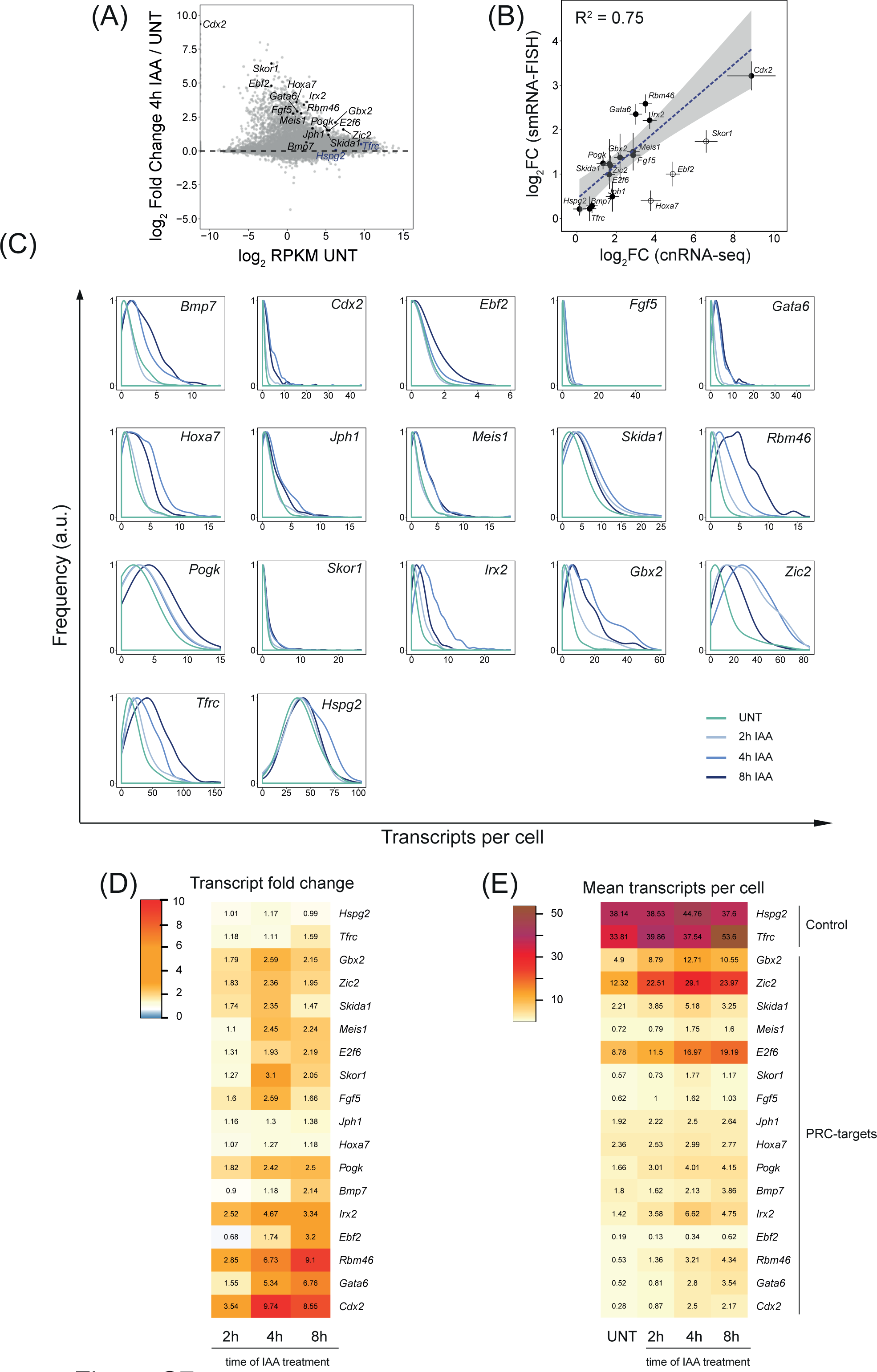
A) An MA plot depicting gene expression (cnRNA-seq) changes following 4h IAA treatment in PRC1^deg^ cells with candidate genes for smRNA-FISH highlighted. The genes chosen span a wide range of initial transcript levels and transcript increase upon RING1B removal. The control genes are highlighted in blue and Polycomb target genes in black. B) Correlation of log2 fold changes between cnRNA-seq and smRNA-FISH after 4 hours of IAA treatment. Hollow dots represent genes excluded from the linear fit. Error bars represent standard error of 3 biological replicates. C) Normalised density plots of transcripts per cells over the time course of RING1B removal for all genes studied with smRNA-FISH. Shown is a representative biological replicate of 3 independent experiments. D) A heatmap of mean fold changes in transcripts per cell over the time course of RING1B removal. Numbers represent the mean of 3 biological replicates. E) As in (D) but representing the mean number of transcripts per cell.

**Supplementary Figure 8.**
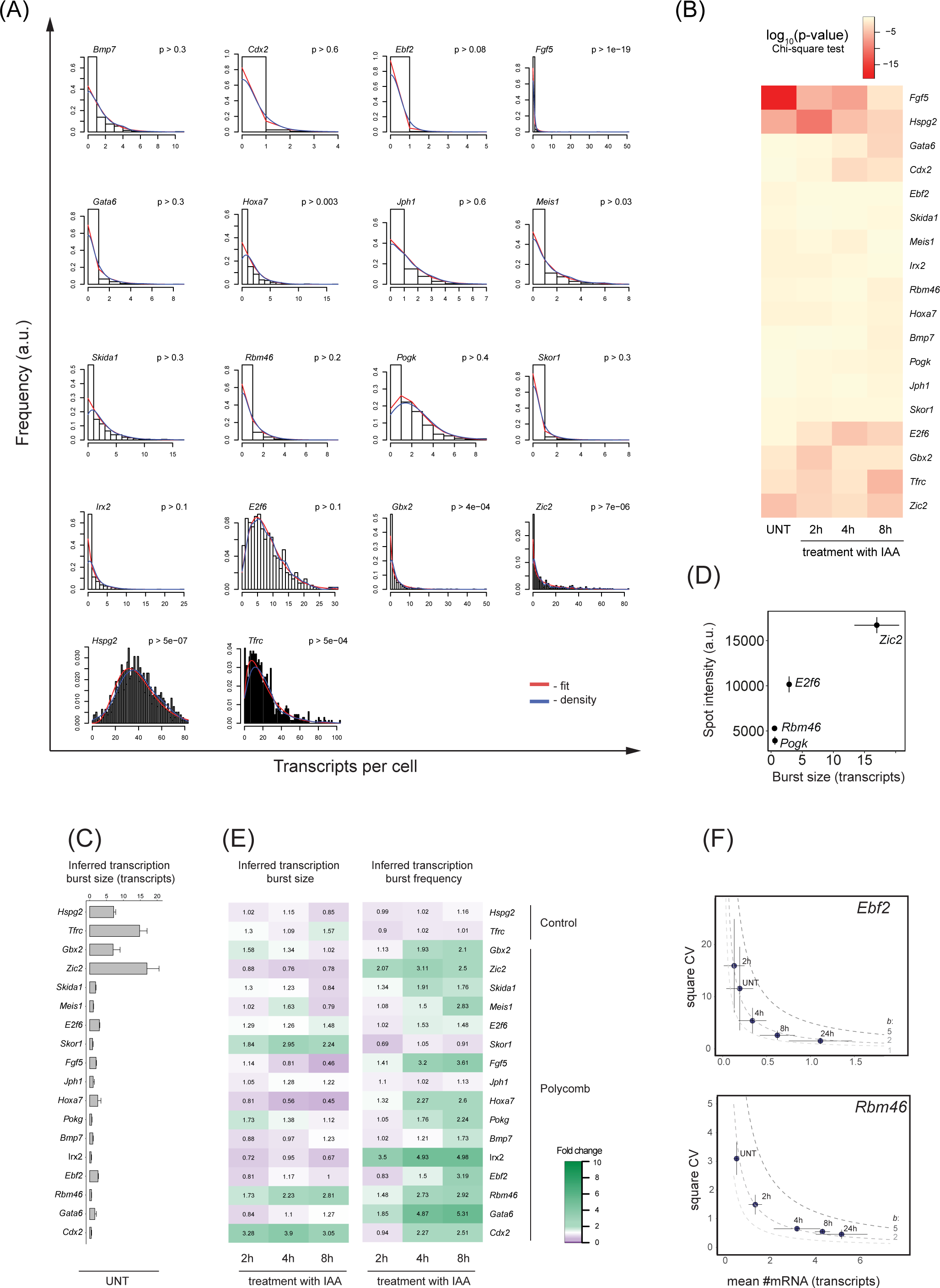
A) Examples of model fits to cellular transcript distributions for all the genes examined. The model fit is in red and density in blue. B) A heatmap of the goodness of fit p-value (Chi-square test) throughout the time course of RING1B depletion. High p-value (light yellow) represents a good negative binomial fit to the data. C) Barplots representing mean inferred transcription burst sizes for Polycomb target genes and control genes. Error bars correspond to standard error of n=3 biological replicates. D) The relationship between nascent spot intensity (active transcription site, measured using nascent smRNA-FISH targeting intronic sequences) and transcription burst size inferred from mRNA-FISH (targeting exonic sequences) reveals that genes with higher predicted burst size values have greater nascent spot intensity. Error bars correspond to standard deviation of n=3 biological replicates. E) Heatmaps illustrating the fold change in inferred transcription burst size (left) and burst frequency (right) over the time course of RING1B depletion for the panel of genes studied. F) Examples of the relationship between square coefficient of variation (CV) and mean number of transcripts per cell demonstrate that Polycomb target genes derepressed upon PRC1 removal experience increase in transcript number while simultaneously retaining constant transcription burst size. Dashed lines represent theoretical relation between CV^2^ and the mean number of transcripts at constant burst size values (CV^2^=*b*/mean#mRNA) with changing burst frequencies. Error bars correspond to the standard deviation of 3 biological replicates.

**Supplementary Figure 9.**
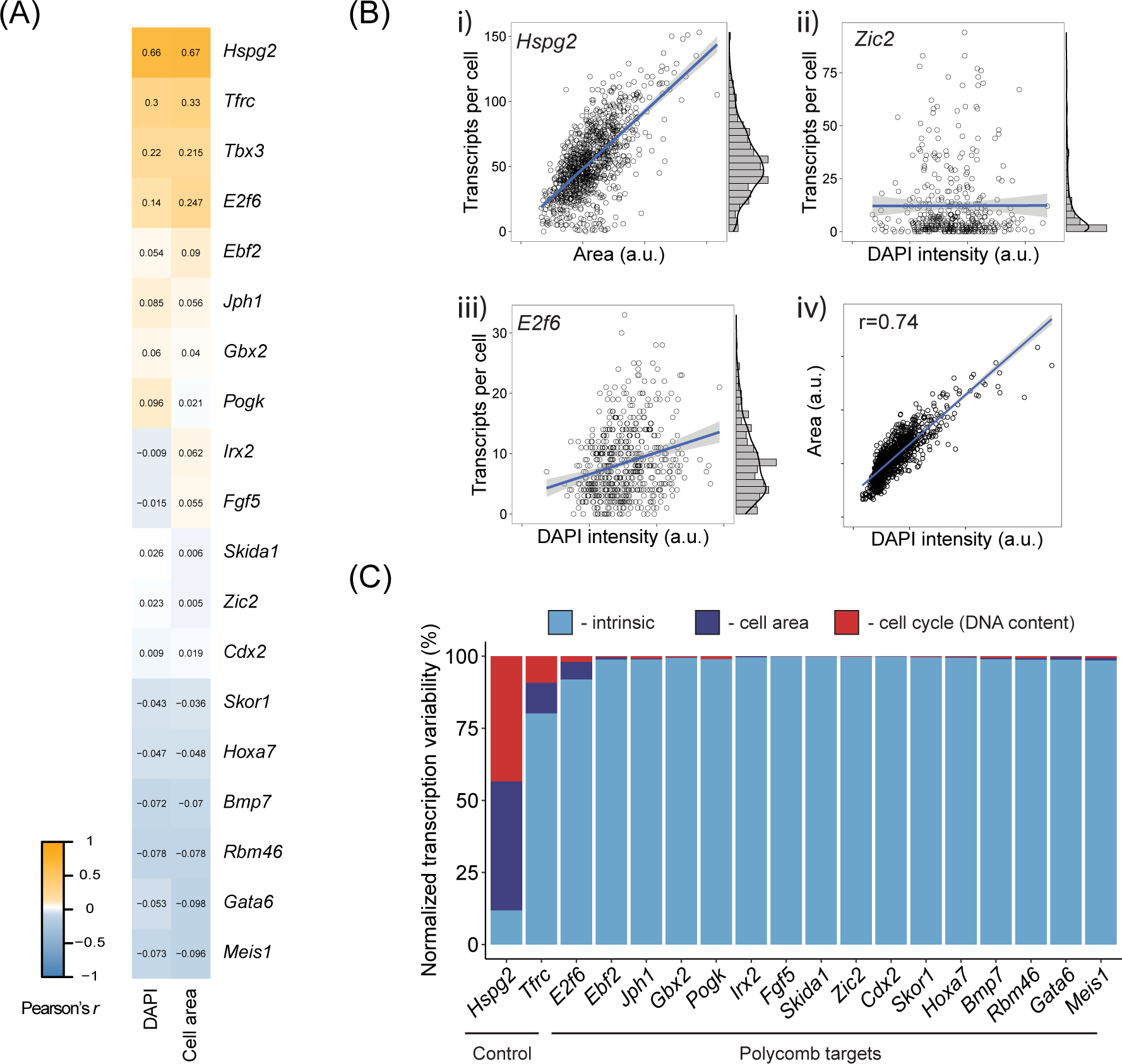
A) Pearson’s correlation coefficient between number of transcripts per cell and either DAPI intensity per cell (a proxy for the cell cycle stage as G2 cells have generally 2x the genetic material of the cells in G1), or cell area (representing cell volume) – the two primary sources of extrinsic noise. B) i – iii) Examples of area or DAPI signal per cell plotted against transcripts per cell for a control (*Hspg2*) or Polycomb target genes (*Zic2*, *E2f6*). Each data-point represents a single cell measurement. iv) Correlation between cell area and DAPI signal intensity (Pearson’s r = 0.74) suggests that volume of ESCs is strongly related to their cell cycle phase. We note that the cell nucleus in ESCs occupies a significant portion of the total cell volume. C) Specific heterogeneity (noise) in transcripts per cell expressed as % of the total heterogeneity measured as square coefficient of variation of the transcripts per cell distributions. Cell area and DAPI signal intensity contribute very little to the overall variability in number of transcripts per cell in a population for Polycomb target genes.

